# Prediction of amphipathic helix – membrane interactions with Rosetta

**DOI:** 10.1101/2020.06.15.152322

**Authors:** Alican Gulsevin, Jens Meiler

## Abstract

Amphipathic helices have hydrophobic and hydrophilic/charged residues situated on opposite faces of the helix. They can anchor peripheral membrane proteins to the membrane, be attached to integral membrane proteins, or exist as independent peptides. Despite the widespread presence of membrane-interacting amphipathic helices, there is no computational tool within Rosetta to model their interactions with membranes. In order to address this need, we developed the AmphiScan protocol with PyRosetta, which runs a grid search to find the most favorable position of an amphipathic helix with respect to the membrane. The performance of the algorithm was tested in benchmarks with the *RosettaMembrane, ref2015_memb*, and *franklin2019* score functions on six engineered and 44 naturally-occurring amphipathic helices using membrane coordinates from the OPM and PDBTM databases, OREMPRO server, and MD simulations for comparison. The AmphiScan protocol predicted the coordinates of amphipathic helices within less than 3Å of the reference structures and identified membrane-embedded residues with a Matthews Correlation Constant (MCC) of up to 0.57. Overall, AmphiScan stands as fast, accurate, and highly-customizable protocol that can be pipelined with other Rosetta and Python applications.

## Introduction

### Definition and importance of amphipathic helices

Amphipathic (or amphiphilic) helices are peptides that have hydrophobic and hydrophilic/charged residues situated on opposite faces of a helix. They can interact with membranes by embedding the hydrophobic residues into the hydrophobic core of the membrane, which leaves hydrophilic/charged residues on the other face interacting with the polar lipid head groups or the solvent. Membrane-interacting amphipathic helices with diverse structure and functions can be found as part of integral and peripheral membrane proteins, and as α-helical peptides with antimicrobial properties (1). The membrane selectivity profile of amphipathic helices may vary based on charge, length, and the amino acid composition of the helix (2) or lipid composition of the membrane (3). Different classes of amphipathic helix motifs have been proposed to fine-tune the behavior of amphipathic helices (4).

In integral and peripheral membrane proteins, helices of amphipathic character play a role in sensing changes at the membrane surface, alteration of membrane curvature, and insertion of proteins into membranes (5). As standalone peptides, positively charged amphipathic helices have been shown to have antimicrobial properties, thus called antimicrobial peptides (AMPs). AMPs disrupt the membrane either by altering the membrane curvature or through formation of pores in the cell membrane (6,7).

### Methods for modeling and analyzing amphipathic helix structures

Despite the widespread presence of amphipathic helices, there are very few methods tailored for modeling of amphipathic helices. 3D-HM and HELIQUEST are servers that take as input a sequence which is modeled as an ideal α-helix, or a user-provided structure file (8,9). They calculate parameters including the magnitude and orientation of the hydrophobic moment, the angle between the hydrophobic moment vector and the membrane normal, and electrostatic potentials of the helices. Images showing the helix orientation with respect to the membrane and electrostatic maps are also provided.

### Monte-Carlo-based methods to model amphipathic helix – membrane interactions

Different than the tools that focus solely on structural modeling of amphipathic helices, the MCPep server utilizes an implicit membrane model with a Monte Carlo approach to predict the position of peptides in the membrane starting with peptide sequences (10,11). The output files include the helical content of the given sequence, trajectory of example simulations, free energy decomposition of the peptide residues, and details on favorable orientations with respect to the membrane. An image file showing the position of the helix with respect to the membrane planes is also provided. One caveat of this method is that the algorithm was not thoroughly benchmarked on amphipathic helices, thus its accuracy is unknown.

### Molecular-dynamics-based modeling of amphipathic helix – membrane interactions

Molecular dynamics (MD) simulations have also been used to model amphipathic helix – membrane interactions of peptides and peripheral membrane proteins (12,13). One advantage of the MD simulations is that the dynamic nature of helix – membrane interactions are reflected in the MD trajectories. Further, the presence of an explicit membrane environment allows observation of specific interaction of amino acids and lipids including changes in the membrane curvature. On the other hand, MD simulations can be time-consuming, resource-intense, and challenging to set up for non-experts.

### Rosetta membrane score functions are fast and accurate alternatives to existing amphipathic helix modeling methods

Considering the caveats of the existing methods, we suggest an alternative approach to obtain fast, accurate, and detailed information on the amphipathic helix - membrane interactions: Rosetta membrane score functions *RosettaMembrane, ref2015_memb,* and *franklin2019* (14–18) are good candidates for evaluating these interactions rapidly considering the success of implicit membrane models in predicting membrane protein structures and coordinates in the past (19,20). The Rosetta membrane score functions *RosettaMembrane* and *franklin2019* are based on the implicit membrane model IMM1 (21) with a default bilayer hydrophobic thickness of 30Å. This membrane model consists of a “hydrophobic core” region of 18Å that represents the non-polar tail groups of lipid molecules and a mid-polar “transition” region of 6Å flanking the hydrophobic core on either side of the membrane that represents the more polar glycerol and carbonyl groups of the lipid molecules. There is no explicit representation of the polar headgroups since these are considered to have the same dielectric constant as the solvent. The *ref2015_memb* score function is based on experimental insertion energies of residues inside the membrane and applies the ‘positive-inside’ rule to Arg, His, and Lys residues.

### AmphiScan is a search algorithm benchmarked on idealized-artificial and naturally-occurring amphipathic peptides

Although the Rosetta membrane energy functions were originally conceived to model integral membrane proteins, we hypothesized that they could be used to score depth and orientations of amphipathic helices as well. Based on this premise, we designed a grid search protocol called AmphiScan that can be customized and pipelined with other Rosetta applications. AmphiScan takes as input a PDB structure and calculates its ideal membrane coordinates based on its score along the membrane. We benchmarked the performance of AmphiScan based on multiple scenarios. First, the algorithm was tested on idealized-artificial α-helices. Six amphipathic helices consisting only of leucine and lysine residues (LK peptides) were modeled as ideal helices and subjected to AmphiScan calculations. Next, five hydrophobic peptides that span the membrane and six hydrophilic peptides from the PDB were subjected to AmphiScan calculations. Finally, 44 amphipathic helix structures belonging to integral and peripheral membrane proteins from the PDB (Supporting Table 1) were tested using the helix membrane coordinates from the OPM, PDBTM, OREMPRO, and MD simulations for comparison. The accuracy of the best energy poses generated by Rosetta was assessed by computing the RMSD from the reference structures and the fraction of residues correctly placed with respect to membrane-embedding.

## Results

### AmphiScan calculations utilize three search parameters to define helix depth, tilt angle, and rotation angle

The AmphiScan protocol was first tested on LK peptides with lengths of 15 to 22 amino acids (22) to assess its success with engineered amphipathic sequences. The six LK peptides LK15, LK18, LK19, LK20, LK21, and LK22 (Table 1) were modeled as ideal α-helices with PyMOL (23). The membrane was placed along the xy-plane with the z-axis as the membrane normal. The rotation angle of the helix around its screw axis was defined as α and the angle of the screw axis with respect to the xy-plane was defined as β (Figure 1). The LK peptides were placed at a β angle of 0° to 180° for independent AmphiScan calculations and α values between 0° and 360° were scored at each β and z value. Multiple step sizes were tested for depth (movement along the z-axis), α, and β parameters. A depth step size of 0.1Å, α angle of 5^0^, and β angle of 1^0^ was selected as a smaller step-size failed to further improve the calculation results but increased the time required. These step-size parameters were used for the naturally-occurring helix sequences as well.

**Table 1:**
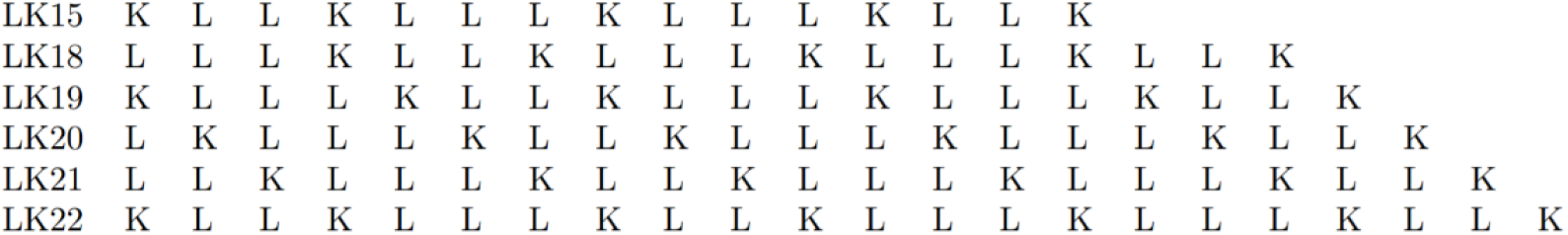
Sequences of the LK peptides used for the AmphiScan calculations.

**Figure 1:**
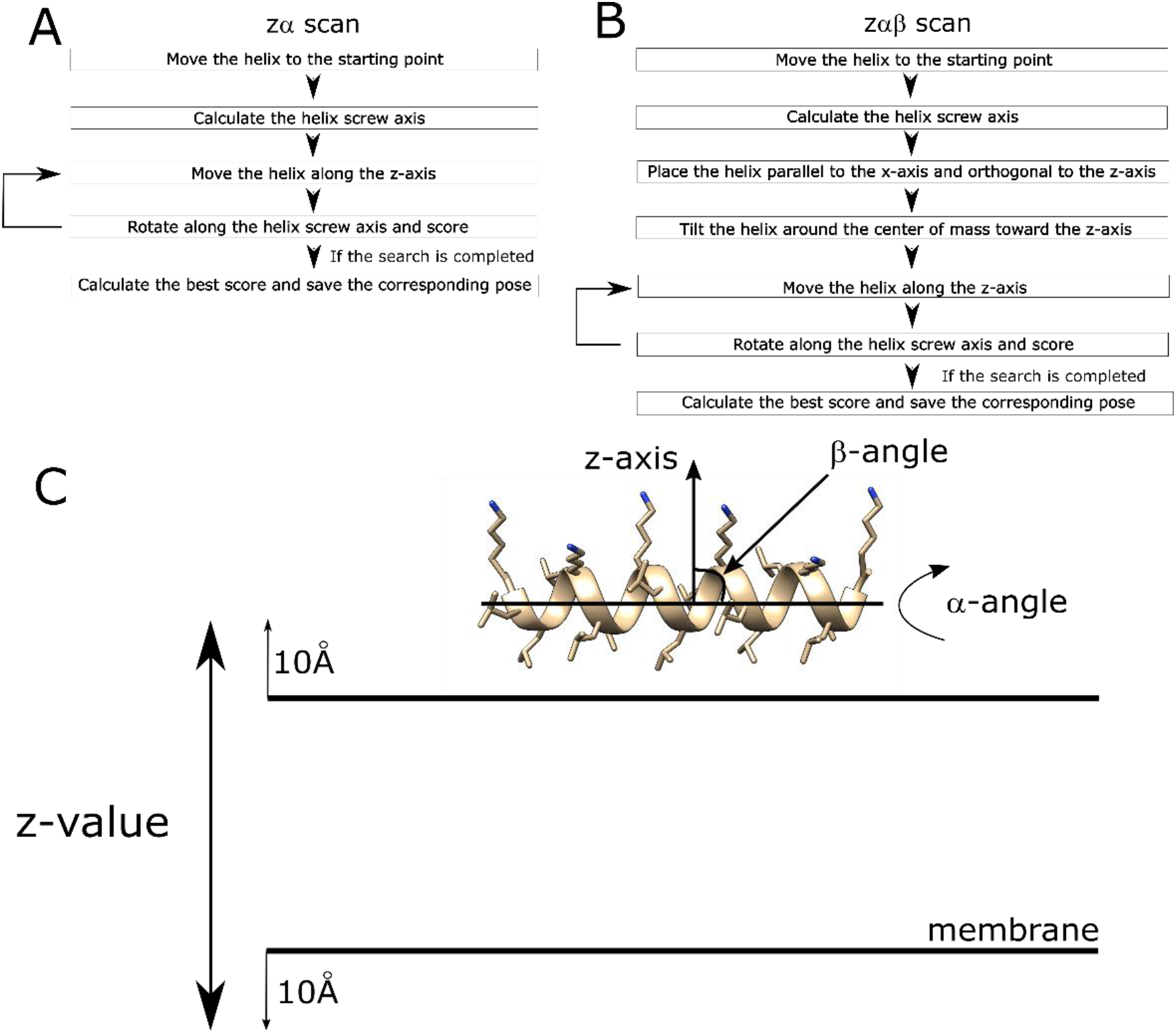
Flowcharts of the AmphiScan calculations: (A) zα-scan flowchart, (B) zαβ-scan flowchart, (C) visual representation of the rotation and tilt operations used to place the helices with respect to the membrane surface. Note that the plane of the membrane corresponds to the xy-plane and the membrane normal corresponds to the z-axis.

### AmphiScan uses a grid search approach to identify the membrane binding configurations of amphipathic helices

Two types of grid searches were run to assess the success of AmphiScan in sampling helix rotation and tilt angles. In the first set of calculations (zα-scans), helix β angles from the reference structures were conserved and only rotation along the helix screw axis and movement along the z-axis was allowed (Figure 1A). In the second set of calculations (zαβ-scans), helix screw axis rotations and helix membrane depth changes were sampled at different β angles, and the pose that gave the best score through all the β angles was kept as the representative pose (Figure 1B). The zα-scans give information about whether the Rosetta membrane score functions can accurately identify the ideal depth and rotation of the α-helix independent of sampling different tilt angles, which is usually the case with amphipathic helices parallel to the membrane surface. The zαβ-scans go a step beyond and give information about whether Rosetta can sample the tilt angle of the helix with respect to the membrane (Figure 1C).

### AmphiScan accurately predicts the membrane depth and orientation of the LK peptides

The zα-scan calculations took 2 – 5 minutes depending on the helix size with the selected parameters on a single CPU. The difference between the best poses from the zα- and zαβ-scans were marginal. For the zαβ-scans of all six helices, the best-scoring pose was obtained at tilt angles of 2 to 6^0^ for all three score functions tested, in accordance with the results of fluorescence spectroscopy and IRRAS experiments that showed near-parallel arrangement of the LK peptides with respect to the membrane surface. Also, the best-scoring structures obtained for all LK peptides were centered 3-4Å below the membrane surface, within the mid-polar region as expected (Supporting Table 2) (24,25). All low-scoring poses had hydrophobic leucine residues embedded in the non-polar core region and the charged ammonium groups of the lysine side chains extending outside the membrane.

### AmphiScan calculations with the *RosettaMembrane* and *franklin2019* score functions illustrate shallow and symmetric energy minima for amphipathic helices

Next, heatmaps color-coded with the scores calculated at different membrane depth and helix rotation angles were plotted for the AmphiScan calculations at a helix arrangement parallel to the membrane surface (e.g. β=0°) to visualize the energy function. In the calculations with the *RosettaMembrane* score function, there were additional low energy regions within the hydrophobic core at the boundary of the mid-polar region. The distribution of the low-energy scores were dispersed at a depth range of ∼1.5Å and a rotation angle range of ∼50^0^ (Figure 2A). The membrane coordinates of the lowest-scoring poses calculated with this score function can be found in Supporting Figure 1, top row.

**Figure 2:**
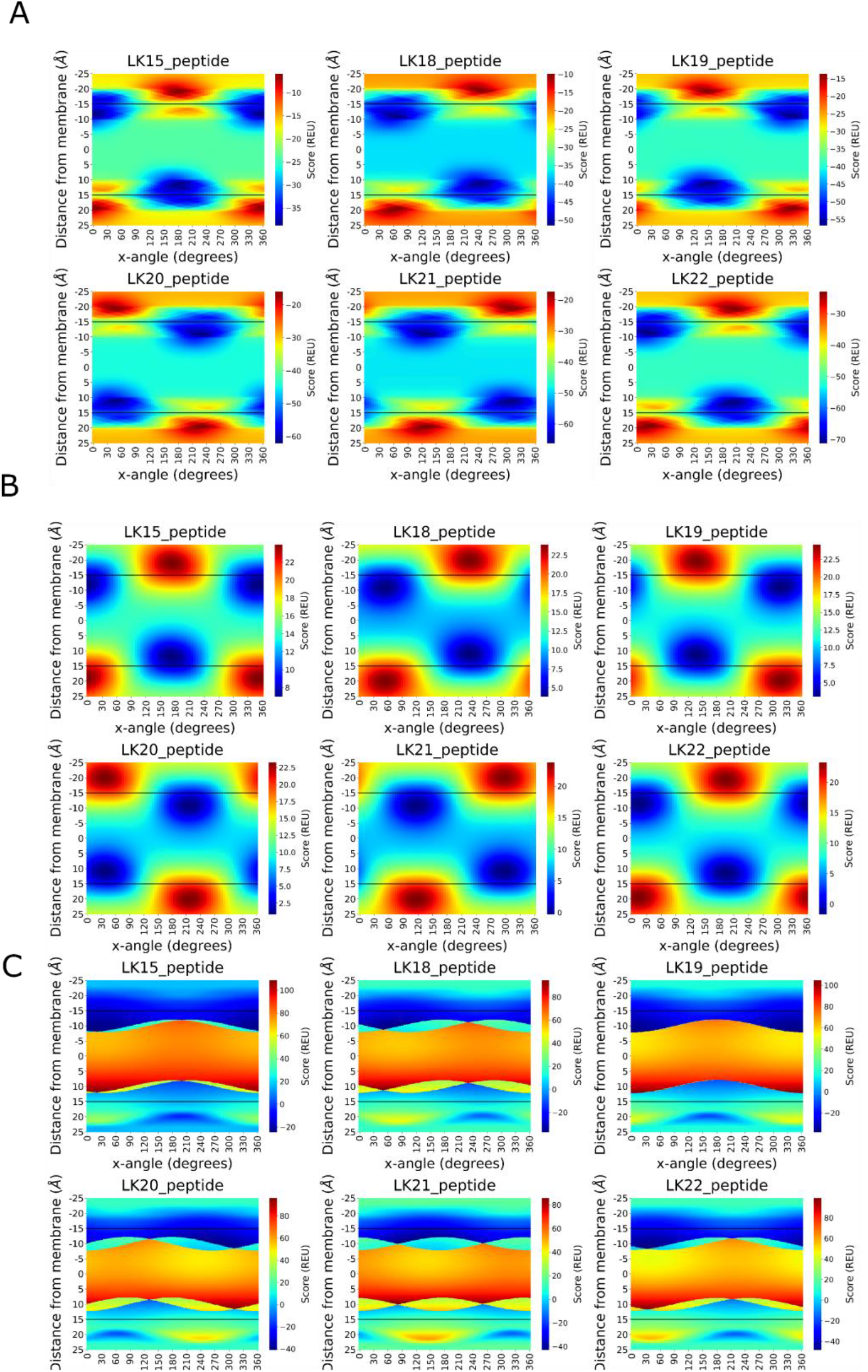
Heatmaps for the Rosetta scans at the best-scoring tilt angle for six LK peptides calculated with the *RosettaMembrane* (A), *franklin2019* (B), and *ref2015_memb* score functions (C). The x-axis represents the rotation along screw axis of the helices and the y-axis stands for the depth of the membrane. Color coding was done separately for each helix where blue color represents the low-scoring regions and the red color represents the high-scoring regions. The black bars represent the boundary of the hydrophobic thickness of the implicit membrane model.

The resulting heatmaps from the *franklin2019* calculations were smoother compared to the *RosettaMembrane* calculations whereby the energy transitions at neighboring grid points were smaller (Figure 2B). However, the low-energy regions incorrectly included large portions of the membrane core, contrary to the *RosettaMembrane* results, where the membrane core displayed the expected higher energies than the low-scoring surface energy wells. These low energy wells were dispersed at a depth range of ∼5Å and a rotation angle of ∼50°. The membrane coordinates of the lowest-scoring poses calculated with this score function can be found in Supporting Figure 1, middle row.

Another expected feature of both heatmaps was the near-symmetric nature of the scores on both sides of the membrane. The regions across the membrane that correspond to a full 180^0^ rotation around the helix screw axis of the lowest-scoring structure gave the lowest scores on the opposite side at the corresponding membrane depth (Figure 2A and 2B). In addition, the α value corresponding to the lowest score yielded the highest scores on the other side of the membrane near the equivalent of the lowest-scoring depth. This is logical since such an arrangement would place the hydrophilic residues inside the hydrophobic core and the hydrophobic residues inside the polar region.

### Ref2015_memb score function is asymmetrical, and insensitive to helix rotation on the positive side

Different than the *RosettaMembrane* and the *franklin2019* score functions, the *ref2015_memb* score function was designed based on the ‘positive-inside rule’, and thus is not symmetric. This asymmetry resulted in a significant difference between the scores obtained on both sides of the membrane (Figure 2C). The scores calculated on one side of the membrane by the algorithm were consistently negative at all rotation angle values, and the lowest-scoring poses were calculated on this side of the membrane (Figure 2C, negative depth values). However, the lowest-scoring poses calculated with this score function had the lysine residues either within the membrane transition region or barely outside the membrane surface contrary to the other two score functions (Supporting Figure 1, bottom row). On the other side, there were narrow low-scoring regions that are ∼5Å outside the membrane. The poses inside the membrane core were heavily penalized by the score function.

### Hydrophobic peptides are placed on the membrane surface or inside the membrane core with inconsistent success for predicting tilt angles

To test whether AmphiScan can also accurately predict the phase placement of non-amphipathic helices, we analyzed hydrophobic peptides that are known to span the membrane and hydrophilic peptides that are soluble. For the membrane-spanning helices, five peptides previously used in implicit membrane benchmarking studies were selected (18,19). The results showed that some peptides have multiple energy minima that may correspond to a positioning on the membrane surface or to a positioning whereby the helices span the membrane with their center of mass placed within the membrane hydrophobic core (Supporting Figure 2)(PDB IDs: 1A11, 2NR1). The deviation from the experimental tilt angles varied from method to method, but no method had a clear superiority over the others in terms of the accuracy of the predicted tilt angles (Supporting Table 3).

### All hydrophilic peptides are placed in the solution phase by the *RosettaMembrane* and *ref2015_memb* score functions but not by *franklin2019*

For the hydrophilic peptide calculations, six water-soluble peptides with available solution NMR structures were selected from the PDB, which are expected to prefer the solvent phase rather than the membrane phase. Consistent with this expectation, all the peptides scored lowest in the solvent region 2.6 to 10Å away from the membrane surface for the *RosettaMembrane* and *ref2015_memb* calculations (Supporting Table 4). Differently, three of the six peptides were placed inside the membrane mid-polar region in the *franklin2019* calculations.

### AmphiScan is benchmarked against membrane coordinates from OPM, PDBTM, OREMPRO, and molecular dynamics simulations

Prediction of the membrane coordinates of naturally occurring amphipathic helices with experimentally determined structures is a more challenging task when compared to engineered systems due to the potentially non-ideal helix pose within the membrane caused by interactions with the other domains of the protein or distorted structure of these helices when using artificial membrane mimetics. There exist myriad experimental methods including NMR, EPR, and fluorescence-quenching to measure membrane immersion depth and orientation with fair accuracy (26). However, systematic analyses of amphipathic helices are rather scant, and the results from these experiments usually depend on environmental conditions and lipid composition of the membrane and thus cannot be directly compared with our results.

Thus, we chose to compare AmphiScan with membrane coordinates predicted by existing computational algorithms trained on experimental data and used these as reference. Naturally, this approach has its flaws as precision and consistency is no guarantee for accuracy. AmphiScan calculations were run with the 44 amphipathic helices in the benchmark set (Supporting Table 1) using the membrane coordinates from the OPM (27–29) and PDBTM (30,31) databases, plus the OREMPRO server (32,33) as references to prevent any biases caused by a single prediction algorithm. Because only the OPM database had membrane coordinates for peripheral membrane proteins and some of the integral membrane proteins in the benchmark set, membrane coordinates approximated from 1μs MD simulations were used as an additional reference point.

### Success of AmphiScan is measured by helix root mean square deviation and membrane-embedded residue Matthews Correlation Coefficient values

Because the membrane coordinates and the identity of the membrane-embedded residues of amphipathic helices play the most important role in helix – membrane interactions, these parameters were used to quantify the error of the AmphiScan calculations. To measure the differences in the membrane depth, orientation, and tilt angles, root mean square deviation (RMSD) values were calculated as the difference between the coordinates of the Cα of each amphipathic helix reference structure from the membrane protein databases (i.e. membrane coordinates) and the best-scoring pose predicted by the AmphiScan protocol. RMSD was selected over individual parameters such as depth, rotation, and tilt angle because it condenses information from all these parameters into a single parameter.

In order to avoid RMSD differences caused by differences between the default Rosetta membrane thickness (30Å) and database-calculated values, the membrane thicknesses calculated by the databases were used as the membrane thickness for each Rosetta calculation. To measure the differences between the identity of the predicted and reference membrane-embedded residue accuracies, Matthews Correlation Constants (MCC) were calculated with respect to the membrane-embedded residues in the reference structures (Methods, Equation 1).

### 44 amphipathic helices from diverse experimental structures were benchmarked with *RosettaMembrane, ref2015_memb, and franklin2019* score functions

We started our analysis using membrane coordinates from the OPM database as the reference for all amphipathic helices. 44 amphipathic helices belonging to 25 peripheral membrane proteins and 19 integral membrane proteins were selected as the benchmark set for the analyses as described in the Materials and Methods section. The benchmark set consisted of amphipathic helices with diverse structural and conformational properties. The helix lengths varied from 8 to 29 amino acids, the tilt angles with the membrane normal were between 0^0^ and 45^0^, and the strength of helix – membrane interactions varied from interacting only with periphery of the membrane surface to being mostly submerged into the hydrophobic core region of the membrane. These 44 structures were subjected to AmphiScan calculations using the *RosettaMembrane, ref2015_memb,* or *franklin2019* score functions.

### AmphiScan calculations with OPM reference structures demonstrate moderate root mean square deviations and good membrane-embedded residue correlations with the *RosettaMembrane* score function

For the zα-scans with the *RosettaMembrane* score function, the average RMSD value was 2.9Å and the average membrane-embedded residue MCC was 0.51. For the zαβ-scans, the average RMSD value was 3.6Å and the average membrane embedded-residue MCC was 0.44 (Supporting Table 5 and Supporting Table 6). The predictive value of the AmphiScan calculations varied among the helices whereby *1h0a_h1, 3j5p_h1, 3jw8_h1, 4hhr_h2, 4zwn_h1, 5ek8_1, 5f19_h4, 5w7b_h1, 5w7l_h1*, and *6d26_h1* had RMSD values below 1Å, but the helices *1q4g_h4, 3tij_h1, 4qnd_h1, 4ymk_h2, 4ymk_h3, 5ahv_h1, 5dqq_h1, 5w7l_h3*, and *6an7_h1* had RMSD values larger than 5Å. Examples to poses with RMSD values smaller than 2Å, between 2 Å and 4Å, and larger than 4Å can be seen in the top, middle, and bottom rows of the Figure 3, respectively.

**Figure 3:**
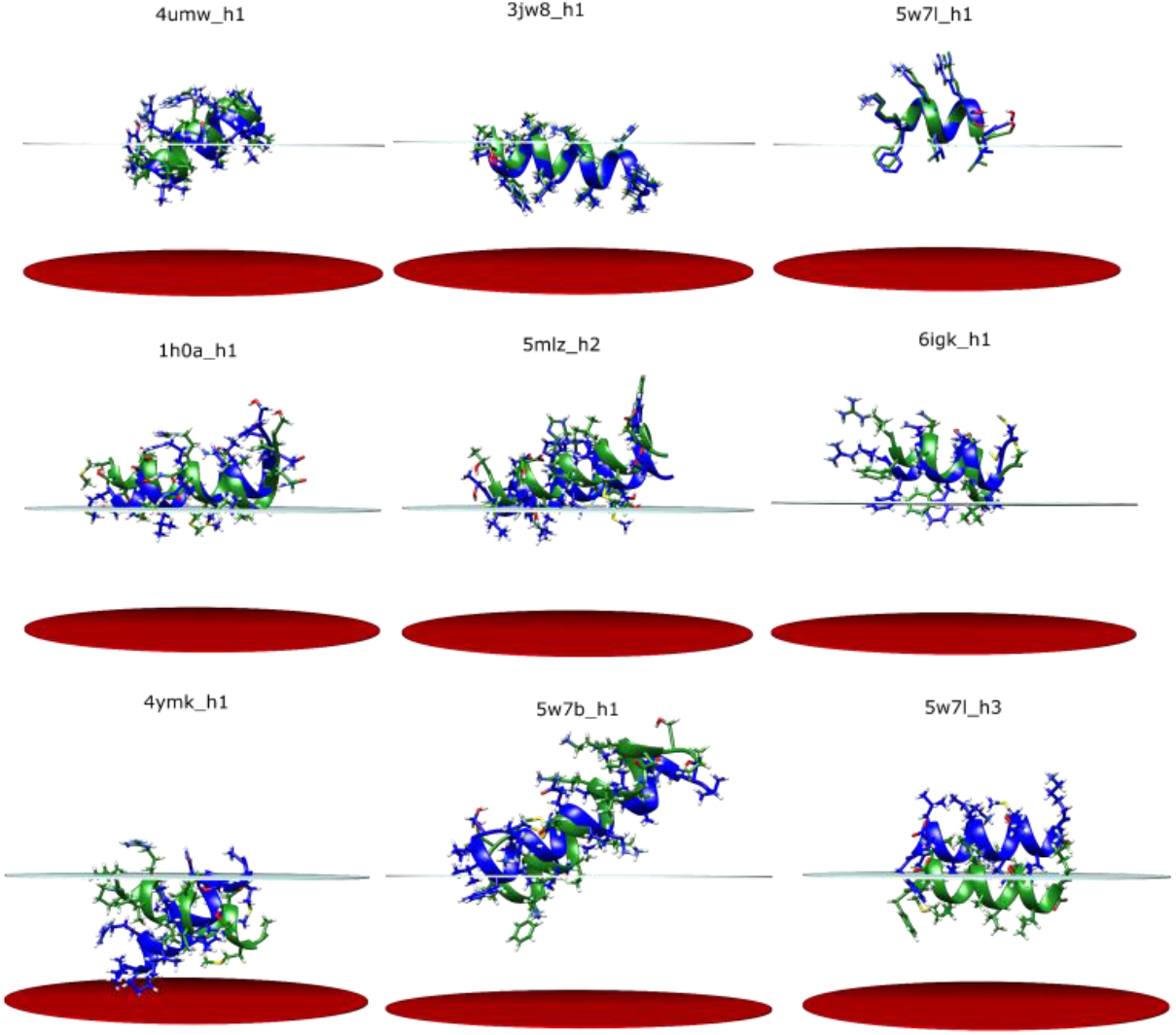
Example alignment poses for the native (green) and best pose structures (blue) from the OPM zαβ calculations. The structures in the top row had RMSD values smaller than 2Å, the middle row had RMSD values between 2 and 4Å, and the bottom row had RMSD values larger than 4Å.

### *Ref2015_memb* and *franklin2019* score functions perform worse than the *RosettaMembrane* score function

The performances of the *ref2015_memb* and *franklin2019* score functions were worse compared to the *RosettaMembrane* score function both in terms of the calculated RMSD values (Supporting Table 5) and the membrane-embedded residue MCC values (Supporting Table 6). For the *ref2015_memb* calculations the RMSD and MCC values were 4.7Å and 0.31, respectively, for the zα calculations and 5.2Å and 0.35 for the zαβ calculations. For the *franklin2019* calculations the RMSD and MCC values were 5.5Å and 0.38 respectively for the zα calculations and 7.0Å and 0.35 for the zαβ calculations.

Visual examination of the poses explained why the performance of *ref2015_memb* and *franklin2019* was worse than the *RosettaMembrane*. The *ref2015_memb* results included both overestimated and underestimated membrane depths with no clear bias towards either case. On the other hand, the depths of the amphipathic helices were frequently overestimated by the *franklin2019* score function placing amphipathic helices in the membrane core region in their entirety (Supporting Table 7). Because *RosettaMembrane* gave consistently better results in the LK peptide, hydrophobic peptide, hydrophilic peptide, and naturally-ocurring amphipathic helix calculations, we only used the *RosettaMembrane* score function for all the following analyses and comparisons with the PDBTM, OREMPRO, and MD results.

### Good recovery of membrane-embedded residues can be obtained at RMSD values as large as 3Å

To understand the relation between the RMSD values and the membrane-embedded residue accuracies, and to determine acceptable RMSD values based on the membrane-embedded residue accuracies, we compared the two values from the *RosettaMembrane* calculations (Supporting Figure 3, left panel). The membrane-embedded residue accuracies were larger than 80% on average below an RMSD of 3Å for both zα- and zαβ-scans with MCC values of 0.50 and 0.57, respectively. At RMSD values larger than 3Å, zα- and zαβ- scans showed different RMSD patterns such that the accuracies decreased steadily at larger RMSD values for the zα- scans, but decreased only slightly for the zαβ-scans. RMSD values lower than 3Å were considered acceptable based on these results since the membrane-embedded residue profiles of the amphipathic helices were accurately represented at RMSD values below this cutoff.

### Side chain repacking has no significant effect on prediction accuracy

In order to assess whether side chain repacking increases the performance of our algorithm, we ran zα calculations with side chain repacking at each rotation step. Helix RMSD and membrane-embedded residue MCC values, and rotamer recovery rates for χ1 and χ2 angles were calculated to measure the success of the algorithm under these conditions. The calculated RMSD (3.0 Å) and membrane-embedded residue MCC values (0.49) were comparable to that of the calculations with no side chain repacking. The rotamer recovery rates measured for the χ1 and χ2 angles were 68% and 41%, respectively (Supporting Table 8). Examples of repacked structures with χ1 rotamer recovery rates over 0.85 can be seen in Supporting Figure 4. Based on lack of decrease in RMSD values or increase in MCC values, we concluded that side chain repacking has no significant effect on the accuracy of the calculations and restricted our analyses to calculations with no side chain repacking.

### AmphiScan performs better with charged helices compared to neutral helices

Following the RMSD and membrane-embedded residue accuracy analyses, we looked for a correlation between the RMSD values and helix properties such as charge, size, number of prolines, experimentally determined tilt angle, native distance from the membrane center, structure quality, and whether the helix is a terminal one to identify biases of the *RosettaMembrane* score function. Of these parameters, helix net charge showed an inverse correlation with the RMSD values, whereby neutral helices on average had the largest RMSD values (Supporting Figure 3, right panel). This correlation was more evident in the zα-scans compared to the zαβ-scans, possibly since fixed helix tilt angles disallow rearrangement of the residues to diminish the unfavorable interactions between the helix and the membrane. The amphipathic helices that were closer to the membrane core in the database structures had a larger ratio of high-RMSD helices compared to peripheral helices, but this effect did not apply to all helices close to the membrane core.

Overall, the average RMSD value calculated for the zα-scans was below our cutoff of 3Å, and the value calculated for the zαβ-scans was 3.6Å. Next, we investigated the effect of database selection on the accuracy of the AmphiScan results.

### Proteins from OPM, PDBTM, and OREMPRO have similar membrane orientations but can have largely different membrane thicknesses

Since amphipathic helices have only tenuous interactions with the membrane surface in general, slight variations in the calculated helix tilt angles or membrane thicknesses predicted by computational algorithms can result in large differences in the identity of the membrane-embedded residues or the apparent immersion depth of amphipathic helices deposited to the databases. In fact, a recent analysis comparing the OPM and PDBTM databases showed that the identity of the predicted membrane-embedded residues in OPM and PDBTM databases match only 60-80% for most proteins (34). Therefore, we used reference structures from the PDBTM database and the OREMPRO server for comparison with the OPM results.

Only 16 of the 44 helices in the benchmark set were available in the PDBTM database. The protein orientations calculated for the proteins hosting these helices were similar in OPM, PDBTM, and OREMPRO with changes in the protein angles less than 5° with respect to the membrane normal. On the other hand, the calculated hydrophobic monolayer thicknesses had differences up to 5Å (Supporting Table 9), which caused drastic differences in the membrane-embedded residue profiles of the amphipathic helices.

### Selection of the reference database significantly affects the root mean square deviation values of individual helices

The functional form of the Rosetta membrane model is independent of membrane thickness, but the RMSD values in this study are calculated based on coordinate differences between the reference structures and AmphiScan poses, which are affected by the differences in membrane thicknesses. To prevent such deviations, membrane thicknesses calculated for each protein by the OPM, PDBTM, and OREMPRO algorithms were used as the hydrophobic thickness of the Rosetta membrane model for the corresponding AmphiScan calculations.

The differences in the calculated hydrophobic thicknesses failed to significantly affect the average RMSD and membrane-embedded residue values, but they significantly affected the outcome of the AmphiScan calculations at the helix level. For both the the zα- and zαβ scans, the average RMSD values calculated with respect to the OPM, OREMPRO, and PDBTM references were similar (Table 2). However, the selected source of the membrane parameters affected the individual RMSD values up to 7.4Å. Also, there was a slight difference between the MCC values of the three methods in both zα (0.40, 0.48, 0.43 for OPM, OREMPRO, and PDBTM) and zαβ (0.35, 0.44, 0.45 for OPM, OREMPRO, and PDBTM) calculations.

**Table 2:** The RMSD values calculated for the best Rosetta zα-scan poses with the *RosettaMembrane* score function where the depth and helix rotation around the screw axis were allowed, and the source of the starting structure used for the calculations. The standard deviations were calculated as the standard deviation among the RMSD values of the same helix with the starting structures from three different sources. All units are in Ångstroms (Å).

| Name | $\alpha$ OPM | $\alpha\beta$ OPM | $\alpha$ OREMPRO | $\alpha\beta$ OREMPRO | $\alpha$ PDBTM | $\alpha\beta$ PDBTM |
| --- | --- | --- | --- | --- | --- | --- |
| 1rhz_h1 | 2.8 | 2.2 | 2.8 | 2.0 | 2.7 | 1.8 |
| 2ziy_h1 | 2.8 | 2.7 | 2.8 | 2.6 | 1.5 | 1.4 |
| 3a7k_h1 | 1.4 | 1.7 | 2.4 | 2.7 | 3.9 | 4.0 |
| 3j5p_h1 | 1.0 | 1.9 | 0.8 | 0.7 | 1.0 | 2.3 |
| 4qnd_h1 | 4.7 | 4.1 | 1.2 | 2.2 | 2.4 | 2.0 |
| 4rp9_h3 | 3.8 | 3.7 | 1.6 | 1.5 | 2.8 | 3.2 |
| 4umw_h1 | 1.6 | 1.7 | 2.1 | 3.2 | 5.1 | 4.5 |
| 4ymk_h1 | 4.2 | 6.7 | 4.1 | 2.0 | 6.2 | 5.4 |
| 4ymk_h2 | 5.4 | 4.4 | 5.3 | 4.3 | 5.6 | 4.1 |
| 4ymk_h3 | 5.0 | 4.5 | 1.7 | 1.7 | 2.4 | 1.6 |
| 5dqq_h1 | 7.2 | 6.8 | 3.0 | 3.2 | 1.6 | 4.6 |
| 5lil_h1 | 3.5 | 6.8 | 1.1 | 7.7 | 1.7 | 4.3 |
| 5mlz_h2 | 2.3 | 2.2 | 5.0 | 2.2 | 1.7 | 2.0 |
| 5uz7_h1 | 2.0 | 1.8 | 9.4 | 9.4 | 4.2 | 8.6 |
| 6an7_h1 | 5.3 | 8.4 | 5.3 | 9.5 | 4.8 | 8.1 |
| 6igk_h1 | 2.1 | 2.0 | 2.0 | 2.0 | 2.8 | 4.4 |
| Average | 3.4 | 3.8 | 3.2 | 3.6 | 3.2 | 3.9 |

### Selection of the lowest root mean square deviation value for each helix improves AmphiScan performance

Because none of the methods used as a reference has a proven accuracy over the other methods, the upper limit of AmphiScan’s success was calculated by considering only the best values for each helix and each reference method. Specifically, the “best possible” RMSD values were calculated by using the RMSD values and the membrane-embedded residue accuracies from the method that gave the best RMSD value for each helix. With this approach, the average RMSD value dropped by around 1.0Å to 2.2Å and the average membrane-embedded residue accuracy MCC value increased from 0.40-0.48 to 0.56 for the zα calculations. For the zαβ calculations, the average RMSD value dropped by 1.1Å – 1.4Å to 2.5Å and the average embedded-residue accuracy MCC value increased from 0.35-0.45 to 0.57, similar to that of the zα-scans. To see whether this improvement also held true for the amphipathic helices with no PDBTM entries, we compared the OPM results with the membrane coordinates calculated by MD simulations.

### Presence of amphipathic helices affects membrane thickness and curvature in MD simulations

We ran 1μs MD simulations in an explicit dioleoyl-phosphatidylcholine (DOPC) membrane model for the 44 amphipathic helices to serve as a second reference point for the AmphiScan calculations. The effect of helix binding on the membrane structure was evident in the MD trajectories. The membrane monolayer thicknesses on the helix-bound side of the membrane were 1.0Å – 1.5Å thinner compared to the other side (Table 3). To include these thinning effects in our calculations, the hydrophobic thicknesses calculated for the helix-bound side of the membrane were used as the monolayer thickness in the AmphiScans rather than half bilayer thicknesses. In addition to membrane thinning, the curvature of the membrane visibly increased around the amphipathic helices, consistent with the distortion analyses on the proteins hosting these helices in the MemProtMD database based on coarse-grain and fully atomistic MD simulations (35).

**Table 3:** The RMSD and membrane-embedded residue accuracies calculated with the *RosettaMembrane* score function in reference to the MD simulations.

| Helix ID | MD thickness | $\alpha$ RMSD | $\alpha$ Stdev | $\alpha$ MCC | $\alpha\beta$ RMSD | $\alpha\beta$ Stdev | $\alpha\beta$ MCC |
| --- | --- | --- | --- | --- | --- | --- | --- |
| 1b4v_h1 | 14.6 | 5.3 | 1.5 | 0.50 | 4.7 | 1.4 | 0.41 |
| 1h0a_h1 | 14.4 | 4.6 | 1.6 | 0.86 | 4.6 | 1.6 | 0.45 |
| 1q4g_h1 | 14.9 | 3.7 | 1.4 | 0.69 | 3.7 | 1.4 | 0.69 |
| 1q4g_h2 | 14.8 | 3.8 | 1.4 | 0.69 | 4.0 | 1.3 | 0.83 |
| 1q4g_h3 | 14.7 | 3.5 | 1.3 | 0.57 | 3.7 | 1.3 | 0.46 |
| 1q4g_h4 | 14.8 | 6.5 | 1.2 | 0.02 | 2.8 | 1.1 | 0.63 |
| 1rhz_h1 | 14.2 | 2.6 | 0.8 | 0.59 | 2.8 | 0.8 | 0.59 |
| 2hih_h1 | 14.4 | 4.8 | 1.2 | 0.55 | 4.2 | 1.2 | 0.66 |
| 2ziy_h1 | 14.8 | 3.4 | 1.4 | 0.76 | 3.5 | 1.4 | 0.76 |
| 3a7k_h1 | 14.8 | 3.9 | 1.5 | 0.41 | 4.6 | 1.5 | 0.50 |
| 3hyw_h1 | 14.2 | 2.9 | 1.0 | 0.38 | 6.9 | 0.8 | -0.15 |
| 3hyw_h2 | 14.6 | 2.6 | 1.0 | 1.00 | 2.6 | 0.9 | 0.82 |
| 3i9v_h1 | 14.6 | 4.1 | 1.4 | 0.60 | 3.7 | 1.3 | 0.77 |
| 3j5p_h1 | 14.6 | 2.6 | 1.0 | 0.70 | 2.7 | 1.0 | 0.54 |
| 3jw8_h1 | 14.7 | 2.0 | 0.9 | 0.35 | 2.0 | 0.9 | 0.35 |
| 3tij_h1 | 13.9 | 3.7 | 0.9 | 0.55 | 2.8 | 0.9 | 0.52 |
| 4hhr_h1 | 14.7 | 3.0 | 0.9 | 0.29 | 2.7 | 0.9 | 0.08 |
| 4hhr_h2 | 14.6 | 3.9 | 1.6 | -0.16 | 2.7 | 1.1 | -0.13 |
| 4hhr_h3 | 14.6 | 8.4 | 1.3 | -0.24 | 2.4 | 0.8 | 0.64 |
| 4m5e_h1 | 14.6 | 4.2 | 1.4 | 0.57 | 4.2 | 1.4 | 0.57 |
| 4nwz_h1 | 14.2 | 2.5 | 1.0 | 0.73 | 6.4 | 1.6 | 0.24 |
| 4qnd_h1 | 14.6 | 3.6 | 1.3 | 0.57 | 2.8 | 1.2 | 0.82 |
| 4rp9_h3 | 14.8 | 4.2 | 1.9 | 0.50 | 4.1 | 1.9 | 0.50 |
| 4umw_h1 | 14.8 | 4.6 | 1.3 | 0.67 | 4.1 | 1.4 | 0.67 |
| 4ymk_h1 | 14.7 | 2.8 | 0.9 | 0.00 | 5.2 | 1.0 | 0.00 |
| 4ymk_h2 | 15.0 | 3.2 | 1.1 | 0.50 | 2.7 | 1.2 | 0.77 |
| 4ymk_h3 | 14.6 | 3.1 | 1.4 | 0.51 | 5.0 | 1.3 | -0.19 |
| 4zwn_h1 | 14.3 | 4.2 | 1.3 | 0.71 | 4.1 | 1.3 | 0.81 |
| 5ahv_h1 | 14.6 | 5.7 | 1.4 | 0.23 | 6.0 | 1.7 | 0.18 |
| 5dqq_h1 | 14.6 | 3.1 | 1.3 | 0.76 | 2.9 | 1.1 | 0.49 |
| 5ek8_h1 | 14.2 | 3.2 | 1.0 | 0.89 | 2.8 | 1.0 | 0.68 |
| 5f19_h3 | 14.7 | 4.6 | 1.6 | 0.67 | 4.5 | 1.6 | 0.67 |
| 5f19_h4 | 14.7 | 7.0 | 2.8 | 0.57 | 3.9 | 1.3 | 0.77 |
| 5lil_h1 | 14.6 | 4.5 | 1.4 | 0.25 | 4.3 | 1.1 | 0.57 |
| 5mlz_h2 | 14.5 | 2.8 | 1.1 | 0.74 | 2.7 | 1.2 | 0.74 |
| 5uz7_h1 | 14.5 | 6.0 | 0.8 | -0.26 | 6.0 | 0.8 | -0.30 |
| 5w7b_h1 | 14.7 | 8.8 | 1.5 | 0.37 | 3.5 | 1.3 | 0.62 |
| 5w7l_h1 | 14.8 | 4.0 | 1.4 | 0.65 | 3.4 | 1.4 | 0.65 |
| 5w7l_h2 | 14.7 | 4.7 | 1.8 | 0.49 | 4.3 | 1.8 | 0.60 |
| 5w7l_h3 | 14.7 | 3.9 | 1.6 | 0.85 | 3.9 | 1.6 | 0.85 |
| 6an7_h1 | 13.8 | 11.1 | 1.5 | 0.00 | 14.4 | 1.4 | 0.00 |
| 6d26_h1 | 14.6 | 3.8 | 1.4 | 0.66 | 3.4 | 1.3 | 0.77 |
| 6dvy_h1 | 13.8 | 8.6 | 1.9 | -0.05 | 7.9 | 2.0 | 0.35 |
| 6igk_h1 | 14.6 | 4.6 | 1.7 | 0.71 | 6.6 | 2.0 | 0.58 |
| Average | 14.6 | 4.4 | 1.3 | 0.49 | 4.2 | 1.3 | 0.50 |

### AmphiScan root mean square deviation values from molecular dynamics simulations are larger than database references though with similar Matthews Correlation Coefficient values

Contrary to the RMSD values calculated using OPM, PDBTM, and OREMPRO membrane coordinates as the reference, the average RMSD values calculated for the zα-scans (4.4Å) were slightly higher than the values calculated for the zαβ-scans (4.2Å). The standard deviations of the RMSD values over 800 frames were between 0.8Å and 2.0Å, depicting the mobile nature of the helices on the membrane surface. Despite the large RMSD difference, the membrane-embedded residue accuracy MCC values were comparable for the zα-scans (0.49) and the zαβ-scans (0.50), higher than the calculations using membrane coordinates from the databases as reference (Table 3). Choosing the best RMSD for each helix from the OPM and MD-simulation reference structures yielded average RMSDs of 2.4Å and 2.9 Å and membrane-embedded residue accuracy MCCs of 0.58 and 0.50 for the zα- and zαβ-scans respectively.

## Discussion

Many integral and peripheral membrane proteins possess amphipathic motifs that may play an important role in protein – membrane interactions. However, structure prediction of these helices on the membrane surface are not trivial. Although a variety of tools exist to model membrane proteins, few methods apply to amphipathic helices that interact with the membrane surface. In this work, we utilized PyRosetta to develop the AmphiScan protocol to predict the ideal depth, orientation, and tilt angle of LK peptides, hydrophobic peptides, hydrophilic peptides, and naturally-ocurring amphipathic helices belonging to peripheral and integral membrane proteins in membranes. The results were analyzed using qualitative experimental data for the ideal peptides and membrane coordinates from OPM, PDBTM, OREMPRO servers and MD simulations for the experimental amphipathic helix structures.

### AmphiScan accurately reproduces membrane coordinates of LK peptides

In the scenario with the engineered LK peptides, the AmphiScan calculations with all three score functions correctly predicted the near-parallel placement of the LK peptides with respect to the membrane surface and the embedding of the helix center of masses to the mid-polar transition region of the membrane. The heatmaps of the scores at different helix screw axis rotation angles and membrane depths showed similar scores within 1.5Å to 5Å of the best-scoring membrane depth and helix rotations up to 50^0^ depending on the selected score function. These variations may be a reflection of the flexible nature of the amphipathic helix – membrane, or the resolution limit of the AmphiScan method.

### Hydrophobic and hydrophilic peptide tilt angle and placement performance varied among the score functions

The performance of all three score functions in placing the hydrophobic peptides in the membrane showed some deviation from the experimental values. Further, some transmembrane helices had multiple low-energy configurations within the membrane, which was previously reported in calculations with these peptides in an implicit membrane model (19). The depth and tilt values calculated in this study with the *ref2015_memb* and *franklin2019* score functions are different than reported in Alford et al. 2020, which showed more consistent results with the experimental data. For the hydrophilic peptides, the *RosettaMembrane* and *ref2015_memb* score functions placed all six peptides into the solution phase, whereas the *franklin2019* score function placed three inside the membrane mid-polar region.

### AmphiScan’s success on naturally occurring helices depend on the source of membrane coordinates, selection of score function, and inclusion of tilt angles

Selection of database had a significant effect on the values calculated for individual helices due to the drastic differences in the hydrophobic membrane thicknesses calculated by the different methods. This suggests that the knowledge of the membrane thickness may improve the quality of the results of the AmphiScan calculations, especially for amphipathic helices that are buried below the mid-polar region.

The score functions had a large effect on the quality of the obtained results, whereby the *ref2015_memb* and *franklin2019* score functions overall showed worse results compared to the *RosettaMembrane* score function, despite the additions made to these score functions to improve their performances. Major issues with these score functions were lack of sensitivity to the scores on one side of the membrane for the *ref2015_memb* score function and overestimation of the amphipathic helix depths for the *franklin2019* score function. Further, shallower placement of the hydrophobic peptides by *RosettaMembrane* suggests a bias of this score function towards placement closer to the membrane surface, which may make it more suitable for calculations with amphipathic helices.

Sampling the tilt angles increases substantially the conformational space that needs to be sampled. Thus, it is not surprising that slightly worse RMSD and MCC values are observed due to the existence of additional low-energy helix placements. For the helices with RMSD values larger than 4Å in the zαβ calculations, this was the major cause of the increased RMSD values.

### AmphiScan’s success is limited by imperfect helix structures and assumptions intrinsic to implicit membrane models

Three main limitations of the AmphiScan protocol are the “non-ideal” structure of the experimental amphipathic helices, the simplistic assumptions behind the *RosettaMembrane* model, and the differences between the reference and Rosetta implicit membrane models.

In theory, an amphipathic helix would be situated on the membrane interface interacting with both the polar and non-polar portions of the membrane, but it may behave differently as part of a larger protein assembly in reality. Several helices were buried inside the hydrophobic core of the membrane despite being identified as amphipathic in the experimental structures. These helices were found to be placed closer to the membrane surface in the zα calculations, which was the most important cause for the large RMSD values observed in these calculations. Another issue was that, most experimental amphipathic helix structures in the benchmark had distorted or kinked structures, which may cause inaccuracies in calculation of helix screw axes, thus perturbing sampling space towards less natural placements. These variations contributed to the larger RMSD values due to minor shifts in the coordinates of the final helix structures. In the future, adding backbone flexibility to these calculations can alleviate this problem.

Other errors are associated with the simplistic nature of the Rosetta membrane models. Properties such as membrane composition, membrane curvature, or disruption of the membrane surface in response to amphipathic helix binding that are known to play a major role in amphipathic helix - membrane interactions (36–39) cannot be fully represented by the implicit membrane models *RosettaMembrane* and *ref2015_memb*. Although the *franklin2019* score function addresses this shortcoming through the use of different membrane types parametrized based on experimental data, overestimation of the helix membrane depths prevented obtaining more accurate results.

Finally, the membrane models used by OPM, PDBTM, OREMPRO, and Rosetta are not identical and treat the non-polar, mid-polar, and polar regions of the membrane differently, which causes ambiguity when comparing the results from different prediction methods.

### Limitations of molecular dynamics simulations make quantification of errors challenging

Although MD simulations are not affected by the limitations mentioned for implicit membrane calculations, helix – membrane interactions in MD simulations may be affected by force field (40,41) or sampling biases (42) leading to helix disruption or unfolding in extreme cases. Also, comparison of membrane coordinates from implicit and explicit membrane calculations may be inaccurate at the membrane boundary since implicit models have no exact physical representation contrary to explicit lipid molecules.

### Future directions

The current algorithm was developed to do a grid search to identify low-scoring regions in the membrane. Although the method was successful in its task, grid searches are computationally expensive. The reason why a grid search was preferred for this algorithm despite the performance limitation was that a full grid search enabled us to extract more information regarding the ideal parameters in a quantitative manner. However, users of the algorithm need not be interested in such detailed information on the system. Therefore, for the future applications of the protocol, a scan algorithm based on gradient minimization will be considered rather than a grid search to reduce the time necessary for the calculations.

## Conclusion

Despite the limitations associated with calculation of helix membrane coordinates and validation of the calculated results, our calculations overall demonstrated that the AmphiScan protocol can predict the embedding depth of amphipathic helices within ∼2.5Å and with a membrane-embedded residue accuracy MCC of ∼0.6 when used with the *RosettaMembrane* score function. Further, the AmphiScan protocol can be coupled with other PyRosetta applications to model ideal α-helices from sequence, which makes it an excellent tool to screen for amphipathic helices based on sequence or structure information.

## Materials and Methods

### Structural modeling of the LK peptides

The LK peptides were selected based on their published amphipathicity and helicity under experimental conditions and the sequences of the highly-helical LK peptides were taken from (24) for the AmphiScan calculations. Each peptide was modeled with PyMOL as an ideal α-helix with φ- and ψ-angles of -57° and -47° respectively.

The *seaborn* module of python was used to plot the heatmaps. The distance and rotation parameters were defined as discrete values and the color coding was done based on the score distribution for the calculation whereby blue color indicates favorable scores and red color indicates unfavorable scores. Color coding was done separately for the score range of each plot. The boundary of the membrane was shown with a black bar at 15Å on both faces.

### Calculations with the hydrophobic peptides

A total of five membrane-spanning hydrophobic peptides were selected peptides were selected based on the benchmark sets used in Ulmschneider et al. 2006 and Alford et al. 2020 (18,19). Five of these structures were downloaded from the PDB (PDB IDs: 1A11, 1PJD, 1PJE, 2NR1) (43–45) and the WALP23 structure was modeled as an ideal α-helix with Rosetta by adjusting the φ and ψ angles of the input sequence to -62 and -41 degrees correspondingly for the sequence GW_2_(LA)_8_LW_2_A (46).

### Calculations with the soluble peptides

Six soluble peptides were selected from the PDB database (PDB IDs: 1FVN, 2B4N, 2KHK, 2MVM, 2O8Z, and 5XNG) based on being dissolved in H_2_O/D_2_O and not including any lipid or detergent molecules (47–52). Each structure was truncated to remove the non-helical regions and was subjected to a relax calculation prior to the AmphiScan calculations (Supporting Table 4). The peptides were placed parallel to the membrane surface and only zα AmphiScan calculations were run since the tilt angles of these peptides are irrelevant. Only membrane depths were calculated for these peptides due to the lack of a reference structure for RMSD calculations.

### Selection of the amphipathic helix structures

Because there is no database for amphipathic helices to select from, ∼3000 peripheral and integral membrane proteins in the OPM database bearing at least one α-helix motif were visually inspected to identify α-helices in close contact with the membrane surface to a total of ∼100 amphipathic helix structures. Only the helices that were explicitly identified as amphipathic by the authors of the primary citation were kept to ascertain the amphipathicity of the selected helices. Amphipathic helices with disulfide bonds to other protein domains and unsolved side chain coordinates were removed from the candidate list to prevent biases caused by conformational restraints in the native structure and side-chain modeling by Rosetta. In addition, the amphipathic helices belonging to protein regions that are known to go through large conformational changes based on literature reports or structures for which differences in the amphipathic helix coordinates were observed in other PDB entries were eliminated. For the remaining 33 protein structures, only the amphipathic helices that are in close contact with the membrane mid-polar region were used for the calculations. When multiple helices were present in a protein structure, the helices were labeled as “*_hn*” where n stands for the nth amphipathic helix within the same protein structure starting from the N-terminus (Supporting Table 1). The final benchmark set consisted of 44 amphipathic helices in total.

### Transformation of the helix coordinates prior to AmphiScan calculations

PyRosetta (53) was used to create all the Rosetta protocols and the RosettaMP framework (54) was used to carry all the membrane protein operations. A new membrane mover was created for the *RosettaMembrane* and *ref2015_memb* calculations based on the RosettaMP *AddMembraneMover* by removing the requirement to load span files since the amphipathic helices usually do not span the membrane. For the *franklin2019* calculations, the *AddMembraneMover* was initiated with the keyword ‘single_TM_mode’ to skip the need for the span file. The membrane was kept fixed centered at (0, 0, 0) and only the protein was allowed to move for the scan calculations. The helices were moved to the scan starting point such that the helix center of mass defined as the centroid of all Cα s was 10Å from the membrane surface along the membrane normal (z-axis).

For the 44 amphipathic helices in the benchmark set, membrane thickness calculated for the whole protein hosting the amphipathic helix was used for each calculation with different reference structures to prevent RMSD variations caused by the differences between the default hydrophobic thickness of *RosettaMembrane* (30Å) and the hydrophobic thicknesses calculated by the databases used to obtain the reference structures. For the LK peptides, hydrophobic peptides, and the hydrophilic peptides the default bilayer thickness was used. All thickness values were rounded to the nearest integer.

### Adjustment of the helix membrane orientations

Following the placement of the helices to the starting z-values, the helices were aligned with the xy-plane for sake of consistency. The helix screw axis was calculated as the difference vector between the average Cα coordinates of the four residues on each end skipping the first and the last residues. The angle between the helix screw axis and x-, y-, and z-axes of the membrane was back-calculated using the definition of the dot product with the following equation:

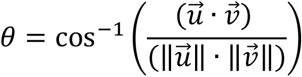

where u stands for the helix screw axis, v stands for the corresponding x-, y-, or z-axis, and θ stands for the angle between the helix screw axis and the coordinate system axis.

For the zα-scans, the tilt angle of the helices from the database coordinates were kept as the angle with the membrane normal, and the helices were only rotated to be orthogonal to the y-axis to retain the original tilt angle. For the zαβ-scans, the helices were first rotated to be orthogonal to the z-axis and then orthogonal to the y-axis to keep the helix screw axis parallel to the x-axis (

Figure 1C). All the geometric operations were done using the *WholeBodyTranslationMover* and *WholeBodyRotationMover* movers of the RosettaMP framework.

### AmphiScan search protocol

Grid scans were run following the alignment operations as summarized in Figure 1. Preliminary calculations with different helix rotation, tilt angle, and depth intervals showed the best resolution of the scores within the shortest amount of time was achieved with a depth interval of 0.1Å, helix rotation angle of 5^°^, and tilt angle of 1^°^, so these values were used as the step sizes for the AmphiScan calculations. Scoring was done with the *mpframework2012* score function, which is the implementation of the *RosettaMembrane* score function to the RosettaMP framework, *ref2015_memb* score function, and *franklin2019* score function. Helices were scored within 10Å from the membrane surface on both ends of the membrane. At each depth value, helices were rotated along their screw axis from 0^°^ to 180^°^. For the zαβ-scans, the helix angle with the y-axis was varied between 0^°^ to 90^°^, and independent AmphiScan calculations were run at each tilt angle. Of the resulting energy profiles, the tilt angle that gave the best-score among all the tilt angles was considered to be the best, and the depth and helix rotation angle values belonging to this tilt angle were considered to be the best parameters for each helix.

### Root mean square deviation and membrane-embedded residue accuracy calculations

RMSD values were calculated as the average root mean squared distance between the coordinates of the helix Cα of the best-scoring Rosetta structure and the corresponding reference structure from OPM, PDBTM, OREMPRO, or MD simulations. The center of mass of each reference structure was subjected to the same translation and rotation operations as the helices to eliminate translational differences between the reference and Rosetta coordinates.

A membrane-embedded residue was defined as a residue with an atom at least 1Å below the membrane surface. Membrane-embedded residue accuracies were calculated according to the following equation:

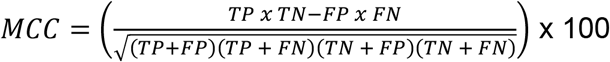

where TP stands for the number of correctly predicted membrane-embedded residues, TN stands for the number of correctly predicted non-membrane-embedded residues, FP stands for the falsely predicted membrane-embedded residues, and FN stands for falsely predicted non-membrane-embedded residues.

### Rotamer recovery rate calculations

Rotamer recovery rate calculations were done using an in-house script utilizing BioPython. Specifically, the input structures used for the AmphiScan calculations were used as the native structures for the comparisons and the lowest-scoring repacked structure was used as the decoy structure. The χ1 and χ2 angles of each residue of the decoy structures were compared with that of the native structures, and residues with rotamer matches within 25° were considered to be recovered.

### Molecular dynamics simulations

The OPM membrane coordinates were used as the reference point for the MD simulations. Explicit DOPC molecules, neutralizing Na^+^ and Cl^-^ ions, and TIP3P water molecules were placed using the CHARMM-GUI server (55). N-terminus acylation (ACE) and C-terminus amidation (NME) was applied to the peptides at the parametrization stage. Each system had a total of ∼100-110 DOPC molecules. The structures from CHARMM-GUI were converted to Amber structure format using a combination of the *charmmlipid2amber.py* script and in-house bash scripts. The DOPC molecules were parametrized with the Amber lipid14 force field (56) and the helix residues were parametrized with the Amber ff14SB force field (57).

The system was minimized in three steps. A restraint weight of 10 kcal/mol Å^2^ was applied to all non-hydrogen atoms in the first step, a restraint weight of 5 kcal/mol Å^2^ was applied to protein backbone (CA, C, O, N) atoms and lipid molecules in the second step, and an unrestrained minimization was run in the third step. The final structure from the third minimization step was heated in four steps, each 5 ns (50K, 100K, 200K, 300K) to a final temperature of 300K in an NVT ensemble with Langevin thermostat. A restraint weight of 5 kcal/mol Å^2^ was applied to the protein backbone CA, C, O, N atoms, and lipid molecules. For each simulation, the system was heated for 2.5 ns and equilibrated at the target temperature for 2.5 ns. The heated system was equilibrated in an NPT ensemble in two 5 ns steps, first with a restraint of 5 kcal/mol Å^2^ on backbone CA, C, O, N atoms and lipid molecules, then with a restraint of 2 kcal/mol Å^2^. Lastly, a 1 μs production run was run in an NPT environment with. At this step of the MD simulations, restraints were applied to the backbone hydrogen bonds to keep the structural integrity of the helices in restrained MD simulations. Specifically, the distance between the carbonyl oxygen atom and the amide hydrogen atoms were calculated for the native structures and the corresponding distances were restrained with a force constant of 10 kcal/mol Å^2^.

### Molecular dynamics trajectory analysis

Trajectory analyses were done with the *pytraj* module of AmberTools (58,59). 800 snapshots from the 1 μs MD trajectories between 200 – 1000 ns were used for the analyses. The distance between the average coordinates of phosphatidylcholine atoms O21 and O11 and the lipid center of mass was used to calculate the monolayer thicknesses of the membrane. The corresponding monolayer thickness of the helix-bound side of the membrane of was used as the thickness in AmphiScan calculations for all helices. RMSD values for the best-scoring AmphiScan poses were calculated with respect to 800 snapshots and the average RMSD values were reported as the final RMSD values.

## Acknowledgements

The authors would like to thank Dr. Julia Koehler Leman for scientific feedback on the project and Dr. Cristina E. Martina for critical reading of the manuscript. Funding for this work was provided by NIH R01 HL144131 and NIH NIGMS R01 GM080403.

## Supporting Tables

**Supporting Table 1:**
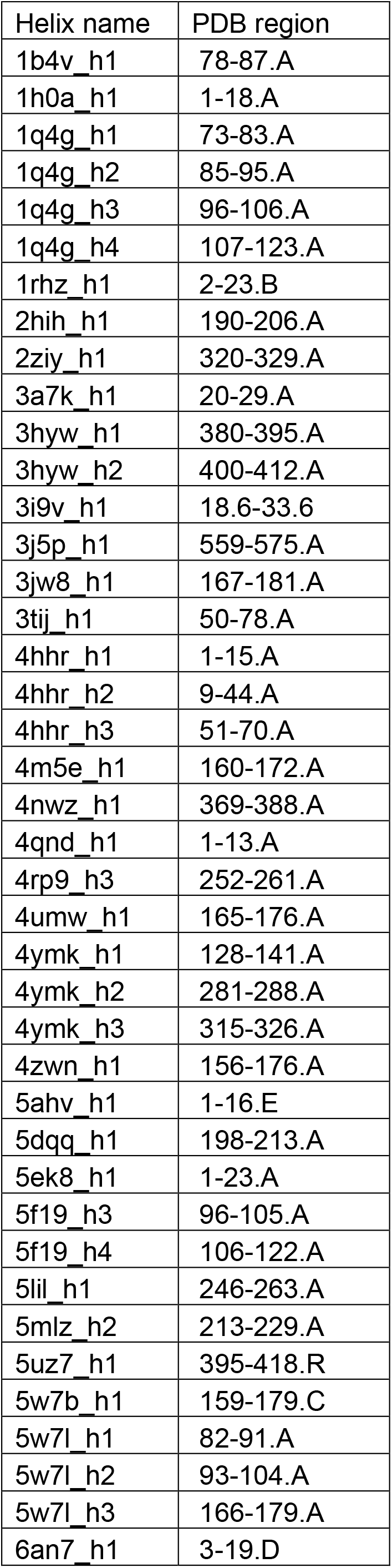
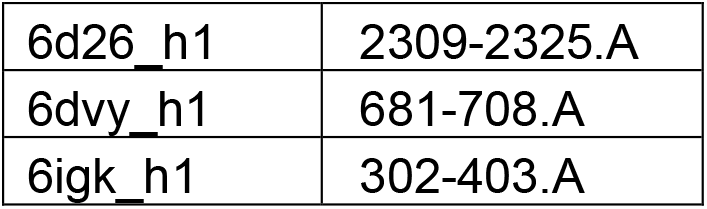
List of the amphipathic helices in the benchmark set and their corresponding residue numbers and chain IDs in the PDB files. The “_xn” denotation was used to distinguish multiple helices belonging to the same protein structure.

**Supporting Table 2:** The depths calculated for the LK peptides using the *RosettaMembrane, ref2015_memb*, and *franklin2019* score functions. Positive depths correspond to helices inside the membrane and negative values indicate helices outside the membrane region. All units are in Angstroms (Å).

| Peptide | RosettaMembrane | ref2015_memb | franklin2019 |
| --- | --- | --- | --- |
| LK15 | 3.4 | 6.9 | 3.1 |
| LK18 | 4.1 | 6.2 | 4.1 |
| LK19 | 3.9 | 7.1 | 3.8 |
| LK20 | 3.8 | 5.1 | 4.2 |
| LK21 | 4.0 | 4.6 | 4.2 |
| LK22 | 3.0 | 6.6 | 3.6 |

**Supporting Table 3:** The tilt angles and depths calculated for 1A11, 1PJD, 1PJE, 2NR1, and WALP23. The tilt angles for the first four peptides were taken from Ulmschneider et al. 2006. The tilt angle for WALP23 was taken from Ozdirekcan et al. 2005. The experimental tilt angles were converted to angle with the membrane plane for consistency. For the helices that had multiple energy minima, the lowest-scoring two were reported as the Tilt angle 1 and the Tilt angle 2. All angles are reported in degrees and all depths are reported in Ångstroms (Å).

| Helix | Experimental tilt angle | Rosetta Membrane Tilt Angle 1 | Rosetta Membrane Tilt Angle 2 | ref2015_memb Tilt Angle 1 | ref2015_memb Tilt Angle 2 | franklin2019 Tilt Angle 1 | franklin2019 Tilt Angle 2 |
| --- | --- | --- | --- | --- | --- | --- | --- |
| 1A11 | 78 | 8 | 90 | 88 | 30 | 86 | N/A |
| 1PJD | 86 | 72 | N/A | 70 | N/A | 54 | N/A |
| 1PJE | 74 | 42 | N/A | 60 | N/A | 14 | 68 |
| 2NR1 | - | 0 | 90 | 66 | 41 | 64 | N/A |
| WALP23 | 82 | 90 | N/A | 34 | 0 | 67 | N/A |

**Supporting Table 4:**
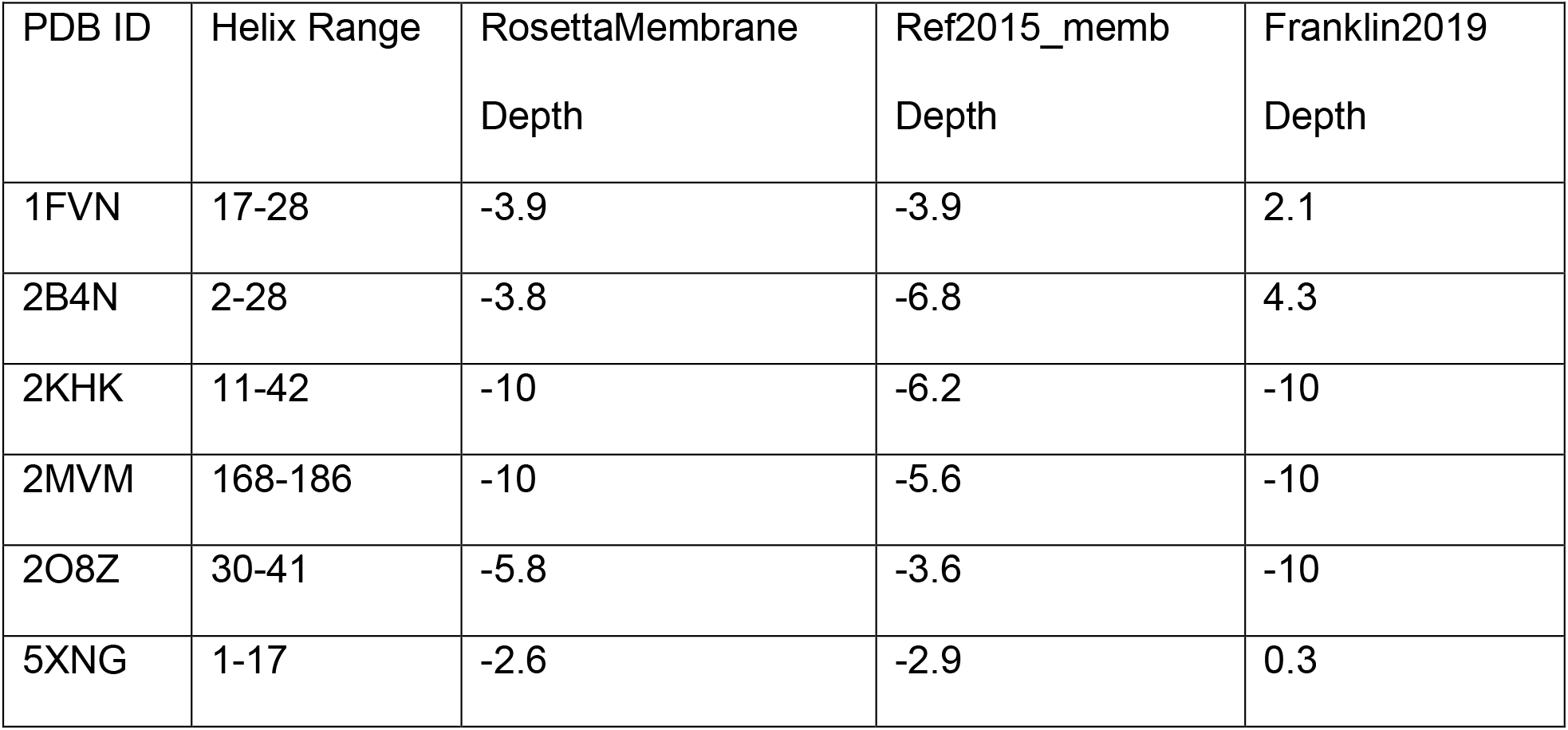
The PDB IDs, helix ranges, and the calculated membrane depths of the tested hydrophilic peptides. Positive depths correspond to helices below the membrane surface and negative values indicate helices outside the membrane region. All units are in Angstroms (Å).

**Supporting Table 5:** The RMSD values calculated for the best Rosetta zα- and zαβ-scan poses with OPM membrane thicknesses using *RosettaMembrane, ref2015_memb*, and *franklin2019* score functions.

| Name | RosettaMembrane RMSD $\alpha$ | ref2015_memb RMSD $\alpha$ | franklin2019 RMSD $\alpha$ | RosettaMembrane RMSD $\alpha\beta$ | ref2015_memb RMSD $\alpha\beta$ | franklin2019 RMSD $\alpha\beta$ |
| --- | --- | --- | --- | --- | --- | --- |
| 1b4v_h<br>1 | 2.1 | 12.0 | 7.1 | 3.9 | 12.0 | 7.2 |
| 1h0a_h<br>1 | 0.5 | 1.7 | 1.7 | 2.2 | 2.1 | 1.6 |
| 1q4g_h<br>1 | 3.6 | 4.8 | 1.2 | 3.6 | 4.8 | 1.4 |
| 1q4g_h<br>2 | 3.2 | 2.7 | 8.3 | 3.7 | 2.2 | 8.8 |
| 1q4g_h<br>3 | 4.2 | 8.0 | 1.8 | 4.2 | 10.2 | 3.5 |
| 1q4g_h<br>4 | 13.3 | 10.3 | 16.1 | 5.7 | 12.1 | 17.0 |
| 1rhz_h<br>1 | 2.8 | 1.7 | 4.7 | 2.2 | 2.2 | 9.4 |
| 2hih_h<br>1 | 1.9 | 2.6 | 6.3 | 1.8 | 2.1 | 6.1 |
| 2ziy_h<br>1 | 2.8 | 3.0 | 2.4 | 2.7 | 2.9 | 6.2 |
| 3a7k_h<br>1 | 1.4 | 2.3 | 4.4 | 1.7 | 1.4 | 6.7 |
| 3hyw_<br>h1 | 3.8 | 3.5 | 8.3 | 5.6 | 3.5 | 14.1 |
| 3hyw_<br>h2 | 2.3 | 5.1 | 5.2 | 2.3 | 4.9 | 5.3 |
| 3i9v_h<br>1 | 1.1 | 2.0 | 4.7 | 2.3 | 1.5 | 6.3 |
| 3j5p_h<br>1 | 1.0 | 5.7 | 3.8 | 1.9 | 11.0 | 4.1 |
| 3jw8_h<br>1 | 0.4 | 4.5 | 0.6 | 0.7 | 5.7 | 3.2 |
| 3tij_h1 | 6.3 | 9.2 | 2.0 | 5.0 | 8.4 | 2.8 |
| 4hhr_h<br>1 | 1.5 | 4.3 | 5.6 | 3.0 | 9.7 | 9.9 |
| 4hhr_h<br>2 | 0.4 | 0.9 | 6.2 | 2.9 | 4.4 | 6.1 |
| 4hhr_h<br>3 | 2.8 | 3.4 | 8.7 | 6.3 | 5.3 | 8.7 |
| 4m5e_<br>h1 | 1.6 | 3.6 | 0.6 | 1.8 | 3.5 | 0.6 |
| 4nwz_<br>h1 | 3.2 | 0.2 | 11.2 | 8.9 | 0.3 | 11.6 |
| 4qnd_h<br>1 | 4.7 | 3.9 | 1.8 | 4.1 | 3.5 | 3.4 |
| 4rp9_h<br>3 | 3.8 | 6.1 | 1.5 | 3.7 | 7.9 | 6.3 |
| 4umw_<br>h1 | 1.7 | 4.0 | 4.3 | 1.7 | 7.0 | 5.0 |
| 4ymk_<br>h1 | 4.2 | 2.1 | 4.6 | 6.7 | 3.0 | 6.0 |
| 4ymk_<br>h2 | 5.4 | 3.8 | 2.1 | 4.4 | 3.4 | 1.6 |
| 4ymk_<br>h3 | 5.0 | 4.9 | 8.4 | 4.5 | 5.0 | 7.7 |
| 4zwn_<br>h1 | 0.9 | 2.8 | 9.6 | 1.7 | 7.4 | 11.8 |
| 5ahv_h<br>1 | 4.7 | 2.4 | 4.1 | 3.5 | 2.7 | 2.5 |
| 5dqq_h<br>1 | 7.2 | 7.1 | 3.2 | 6.8 | 6.9 | 8.3 |
| 5ek8_h<br>1 | 0.3 | 1.7 | 9.0 | 0.7 | 3.1 | 11.0 |
| 5f19_h<br>3 | 1.9 | 6.9 | 1.9 | 1.8 | 7.0 | 1.8 |
| 5f19_h<br>4 | 0.6 | 9.2 | 15.8 | 6.3 | 9.5 | 16.0 |
| 5lil_h1 | 3.5 | 3.2 | 7.5 | 6.8 | 2.4 | 13.2 |
| 5mlz_h<br>2 | 2.3 | 2.3 | 8.6 | 2.2 | 2.4 | 8.2 |
| 5uz7_h<br>1 | 2.0 | 2.0 | 3.9 | 1.8 | 2.0 | 6.3 |
| 5w7b_h1 | 0.5 | 18.4 | 9.7 | 7.3 | 9.1 | 13.1 |
| 5w7l_h1 | 0.9 | 6.9 | 1.6 | 0.8 | 7.6 | 4.8 |
| 5w7l_h2 | 2.1 | 3.4 | 2.2 | 2.0 | 3.4 | 2.6 |
| 5w7l_h3 | 5.5 | 3.1 | 5.4 | 5.5 | 10.5 | 8.4 |
| 6an7_h1 | 5.3 | 5.9 | 7.9 | 8.4 | 6.4 | 8.7 |
| 6d26_h1 | 0.4 | 1.6 | 4.1 | 1.2 | 1.5 | 5.6 |
| 6dvy_h1 | 2.9 | 3.3 | 9.7 | 2.0 | 3.7 | 12.0 |
| 6igk_h1 | 2.1 | 9.5 | 3.1 | 2.0 | 2.7 | 3.1 |
| Average | 2.9 | 4.7 | 5.5 | 3.6 | 5.2 | 7.0 |

**Supporting Table 6:**
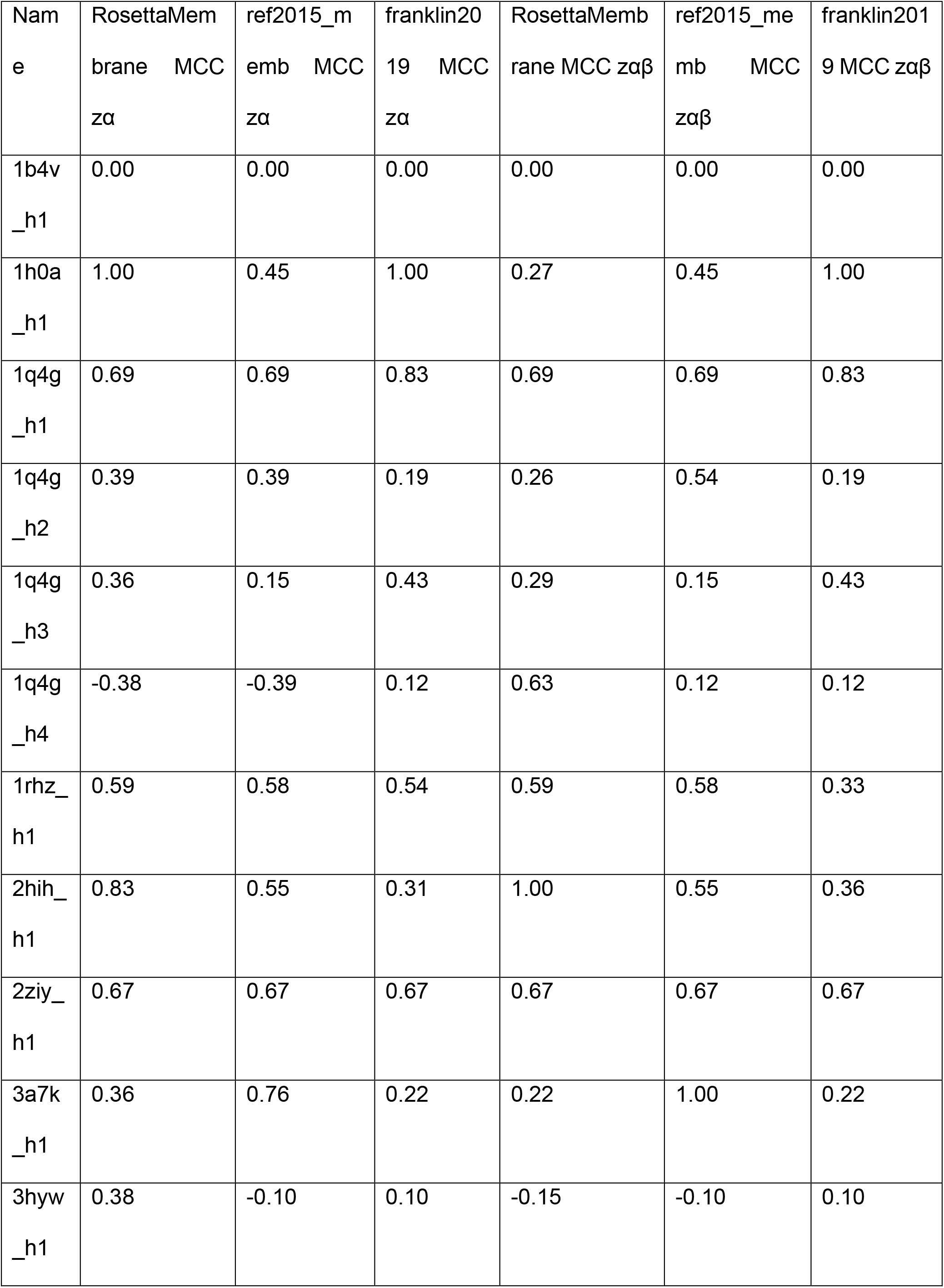
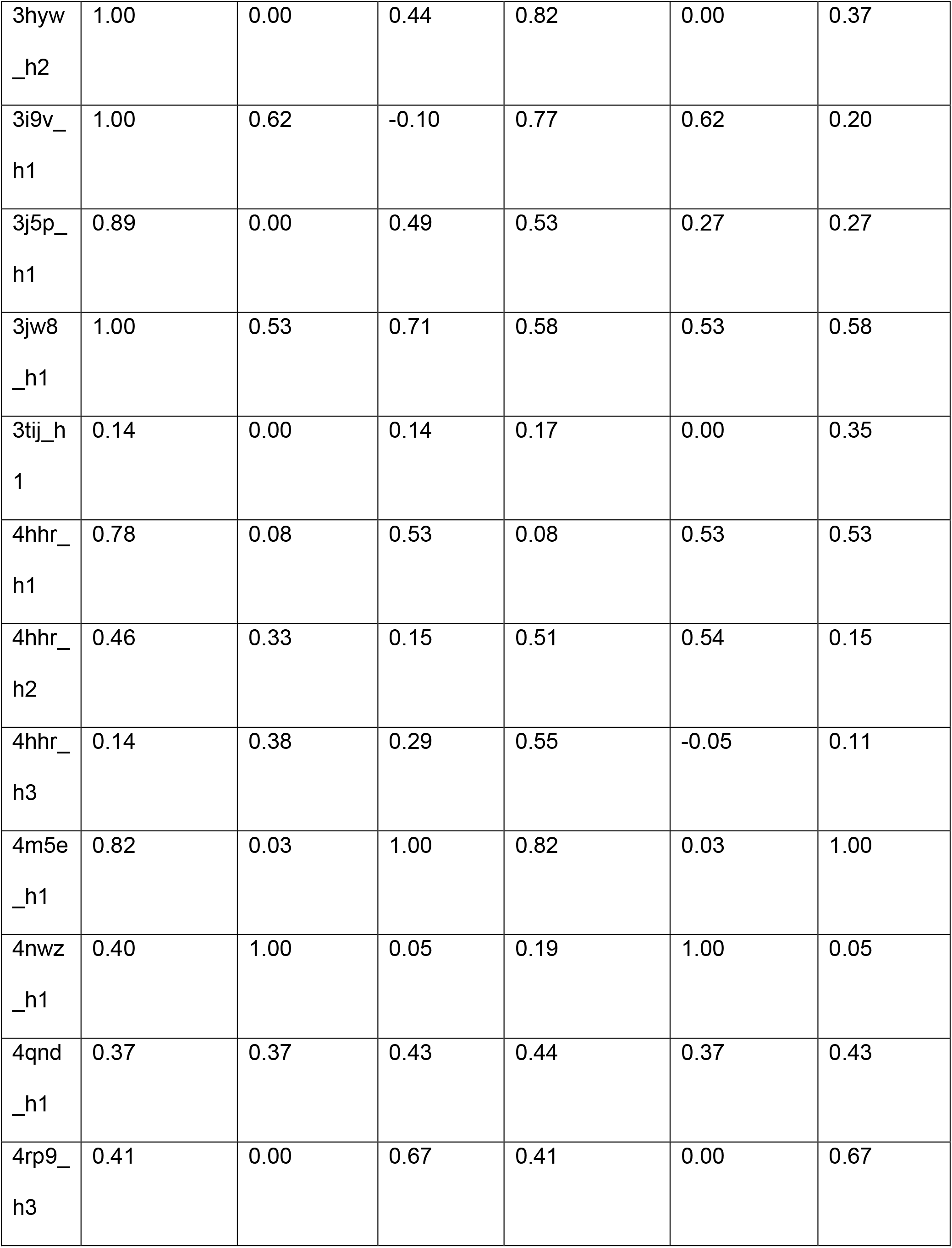
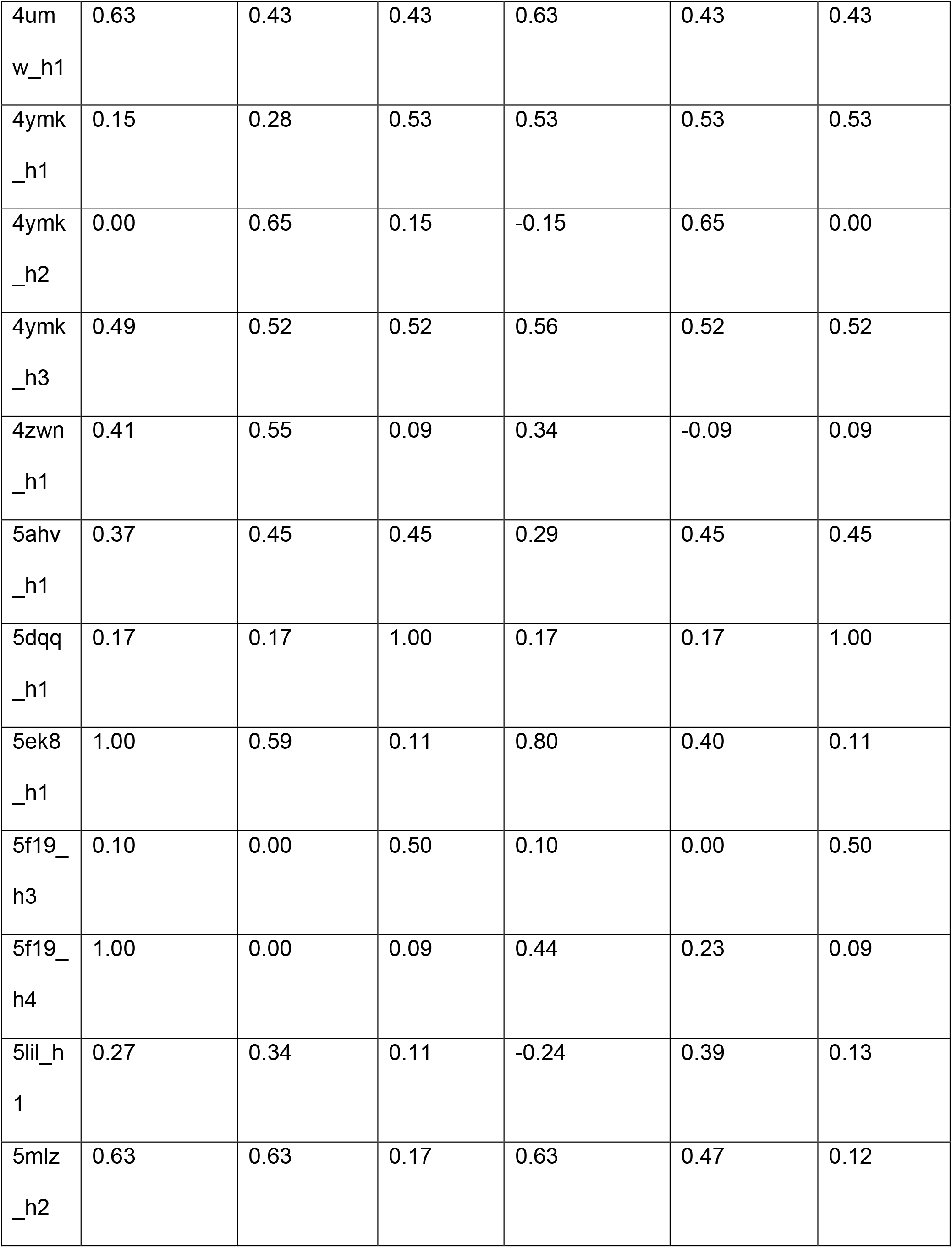
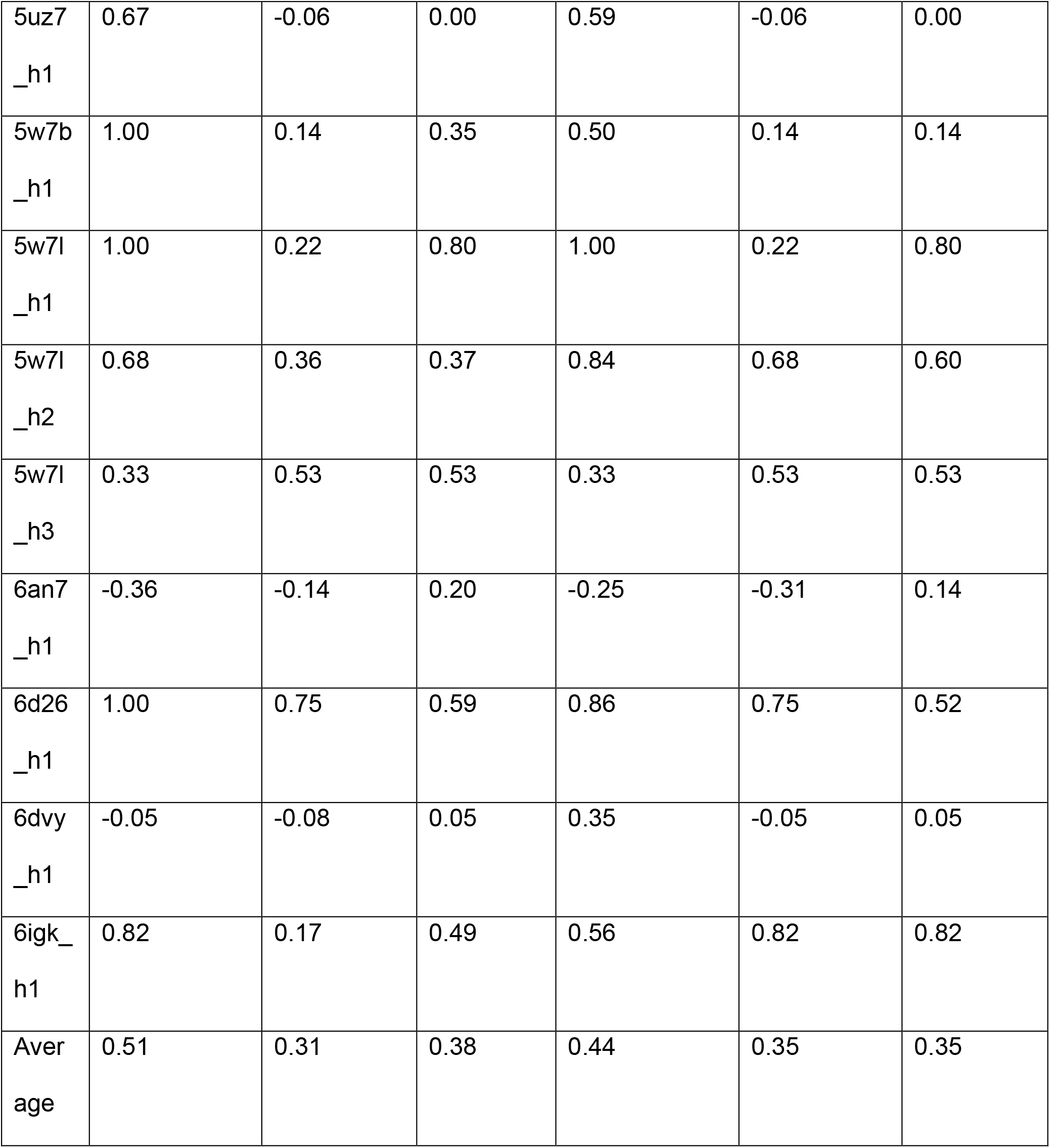
The MCC values calculated for the best Rosetta zα- and zαβ-scan poses with OPM membrane thicknesses using *RosettaMembrane, ref2015*_memb, and *franklin2019* score functions.

**Supporting Table 7:**
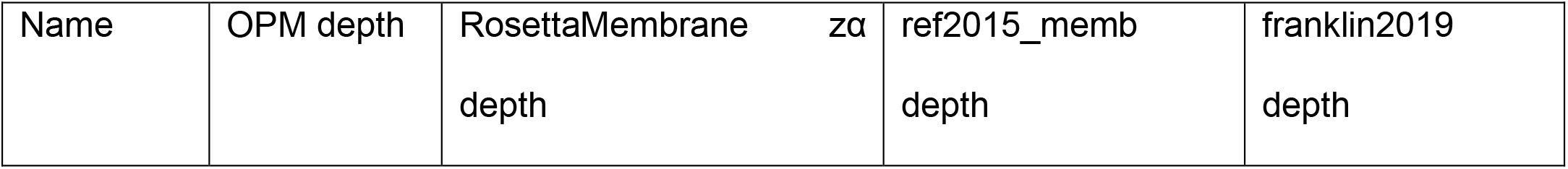
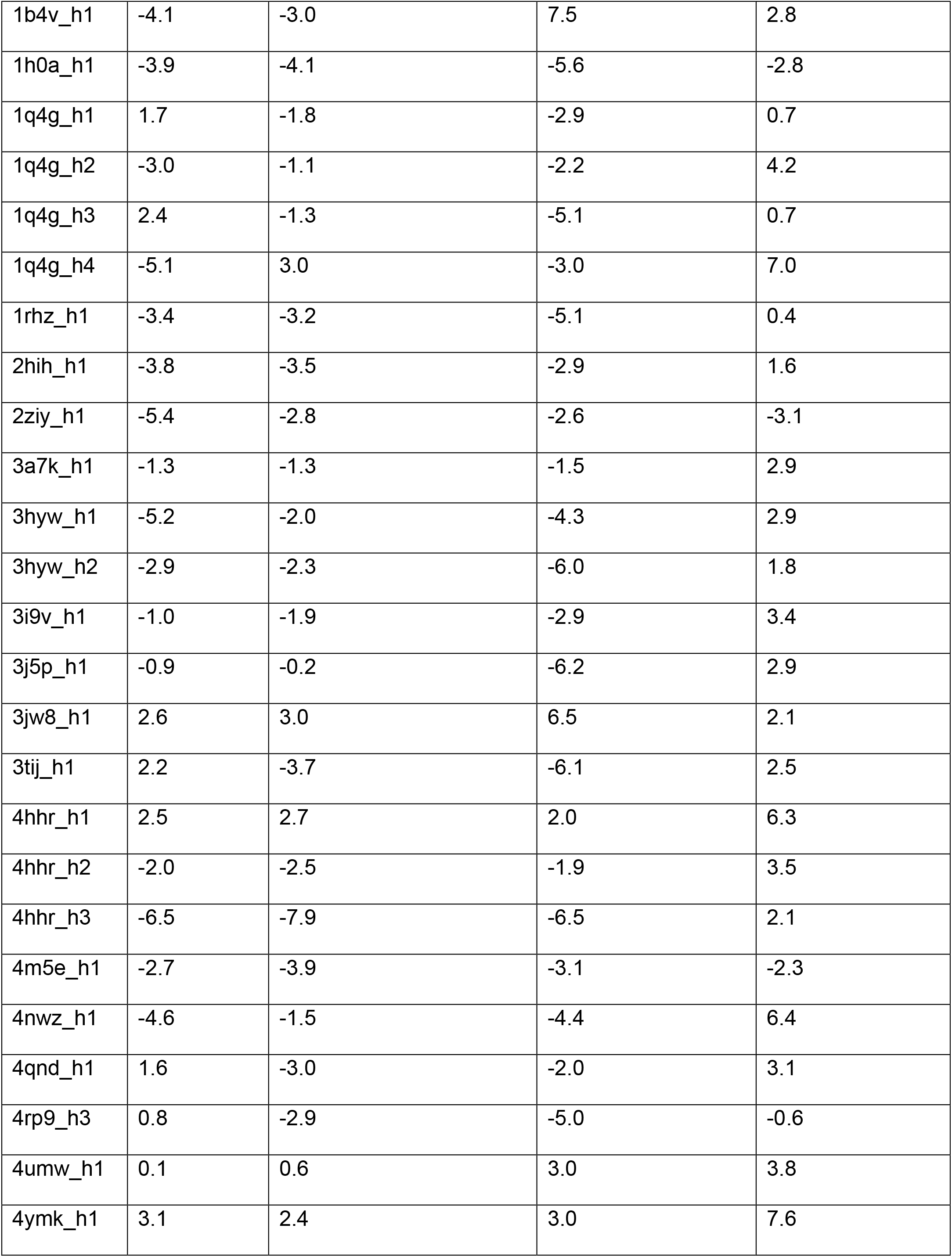
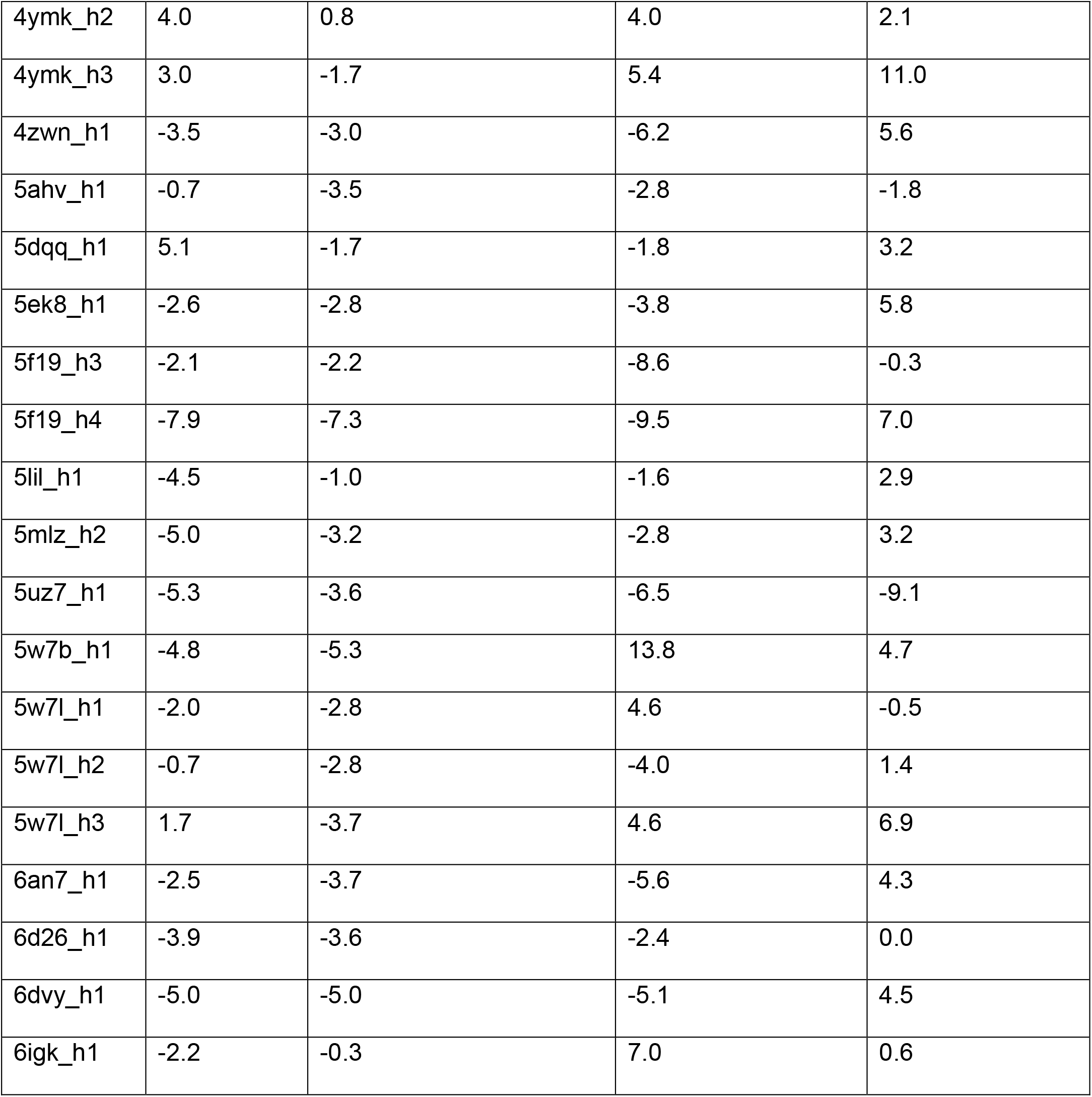
The depths calculated for the OPM structures and the best zα-scan pose calculated with the *RosettaMembrane, ref2015_memb*, and *franklin2019* score functions.

**Supporting Table 8:** The RMSD values, MCC values, and rotamer recovery rates for χ1 and χ2 belonging to the zα calculations with side chain repacking calculated with the *RosettaMembrane* score function.

| Helix name | RMSD (Å) | MCC | Rotamer recovery rate ($\chi_1$) | Rotamer recovery rate ($\chi_2$) |
| --- | --- | --- | --- | --- |
| 1b4v_h1_renum | 2.1 | 0.00 | 0.70 | 0.50 |
| 1h0a_h1_renum | 0.5 | 1.00 | 0.61 | 0.39 |
| 1q4g_h1_renum | 3.6 | 0.69 | 0.64 | 0.46 |
| 1q4g_h2_renum | 3.1 | -0.04 | 0.91 | 0.55 |
| 1q4g_h3_renum | 4.2 | 0.36 | 0.67 | 0.56 |
| 1q4g_h4_renum | 10.2 | -0.26 | 0.53 | 0.24 |
| 1rhz_h1_renum | 2.8 | 1.00 | 0.55 | 0.14 |
| 2hih_h1_renum | 2.0 | 0.83 | 0.50 | 0.19 |
| 2ziy_h1_renum | 2.8 | 1.00 | 0.67 | 0.44 |
| 3a7k_h1_renum | 1.3 | 0.65 | 0.60 | 0.30 |
| 3hyw_h1_renum | 5.0 | -0.17 | 0.73 | 0.33 |
| 3hyw_h2_renum | 2.3 | 0.82 | 0.62 | 0.31 |
| 3i9v_h1_renum | 1.3 | 1.00 | 0.39 | 0.15 |
| 3j5p_h1_renum | 0.9 | 0.76 | 0.47 | 0.27 |
| 3jw8_h1_renum | 0.4 | 1.00 | 0.69 | 0.39 |
| 3tij_h1_renum | 6.1 | 0.14 | 0.63 | 0.38 |
| 4hhr_h1_renum | 1.5 | 0.78 | 0.73 | 0.47 |
| 4hhr_h2_renum | 1.2 | 0.67 | 0.67 | 0.47 |
| 4hhr_h3_renum | 2.9 | 0.14 | 0.79 | 0.58 |
| 4m5e_h1_renum | 1.9 | 0.82 | 0.73 | 0.36 |
| 4nwz_h1_renum | 3.2 | 0.35 | 0.81 | 0.69 |
| 4qnd_h1_renum | 4.6 | 0.37 | 0.80 | 0.60 |
| 4rp9_h3_renum | 3.8 | 0.27 | 1.00 | 0.50 |
| 4umw_h1_renum | 1.6 | 0.63 | 0.60 | 0.50 |
| 4ymk_h1_renum | 4.2 | 0.15 | 0.75 | 0.67 |
| 4ymk_h2_renum | 5.4 | 0.00 | 1.00 | 0.50 |
| 4ymk_h3_renum | 5.0 | 0.41 | 1.00 | 0.70 |
| 4zwn_h1_renum | 0.8 | 0.65 | 0.53 | 0.16 |
| 5ahv_h1_renum | 2.2 | 0.52 | 0.86 | 0.57 |
| 5dqq_h1_renum | 7.2 | 0.17 | 0.33 | 0.13 |
| 5ek8_h1_renum | 1.3 | 1.00 | 0.71 | 0.38 |
| 5f19_h3_renum | 1.9 | 0.10 | 0.89 | 0.56 |
| 5f19_h4_renum | 0.6 | 1.00 | 0.63 | 0.31 |
| 5lil_h1_renum | 3.5 | 0.34 | 0.80 | 0.33 |
| 5mlz_h2_renum | 2.2 | 0.55 | 0.56 | 0.31 |
| 5uz7_h1_renum | 1.8 | 0.34 | 0.68 | 0.27 |
| 5w7b_h1_renum | 11.8 | -0.45 | 0.74 | 0.47 |
| 5w7l_h1_renum | 1.0 | 0.76 | 0.56 | 0.56 |
| 5w7l_h2_renum | 2.2 | 0.84 | 0.55 | 0.27 |
| 5w7l_h3_renum | 5.3 | 0.39 | 0.92 | 0.69 |
| 6an7_h1_renum | 5.3 | -0.31 | 0.38 | 0.06 |
| 6d26_h1_renum | 0.4 | 1.00 | 0.59 | 0.41 |
| 6dvy_h1_renum | 0.3 | 0.46 | 0.59 | 0.41 |
| 6igk_h1_renum | 2.1 | 0.82 | 0.70 | 0.50 |
| Average | 3.0 | 0.49 | 0.68 | 0.41 |

**Supporting Table 9:** Hydrophobic thicknesses calculated by different methods for each protein. The thickness calculations were done on full proteins, and the same thickness value was used for all the helices belonging to the same protein structure. Standard deviations were calculated for the different thicknesses given by different methods for the same protein structure. All units are in Ångstroms (Å).

| PDB ID | OPM | OREMPRO | PDBTM | Stdev |
| --- | --- | --- | --- | --- |
| 1rhz | 14.6 | 14.5 | 14.5 | 0.1 |
| 2ziy | 16.5 | 17.0 | 15.3 | 0.9 |
| 3a7k | 16.9 | 16.0 | 13.8 | 1.6 |
| 3j5p | 14.2 | 16.6 | 16.5 | 1.4 |
| 4qnd | 16.5 | 12.0 | 16.0 | 2.5 |
| 4rp9 | 14.9 | 10.5 | 13.0 | 2.2 |
| 4umw | 15.5 | 11.5 | 12.8 | 2.0 |
| 4ymk | 13.8 | 12.3 | 11.5 | 1.2 |
| 5dqq | 14.9 | 11.1 | 12.5 | 1.9 |
| 5lil | 15.5 | 19.5 | 18.0 | 2.0 |
| 5mlz | 15.3 | 14.3 | 14.8 | 0.5 |
| 5uz7 | 15.1 | 20.0 | 18.0 | 2.5 |
| 6an7 | 14.9 | 15.5 | 14.3 | 0.6 |
| 6igk | 17.1 | 18.0 | 14.8 | 1.7 |

## Supporting Figures

**Supporting Figure 1:**
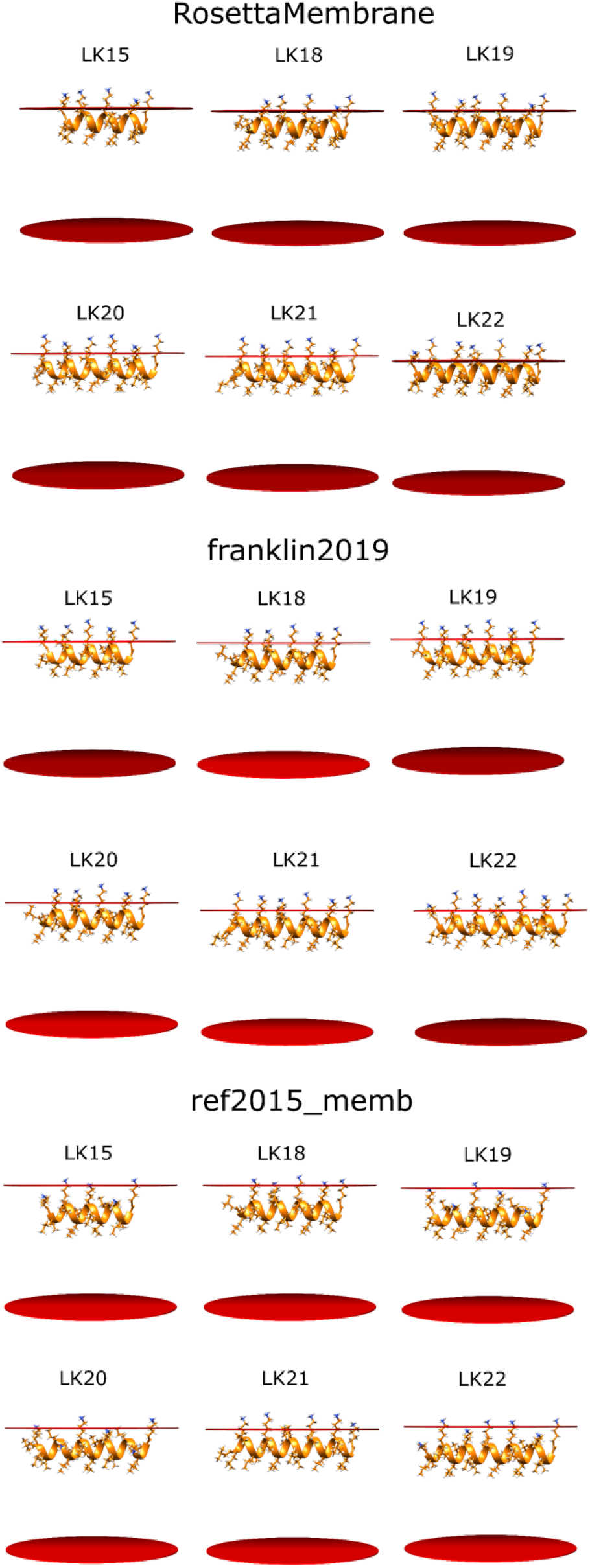
Structures of the lowest-scoring AmphiScan poses o the LK peptides calculated with the *RosettaMembrane, franklin2019*, and *ref2015_memb* score functions. The red planes represent the membrane surface.

**Supporting Figure 2:**
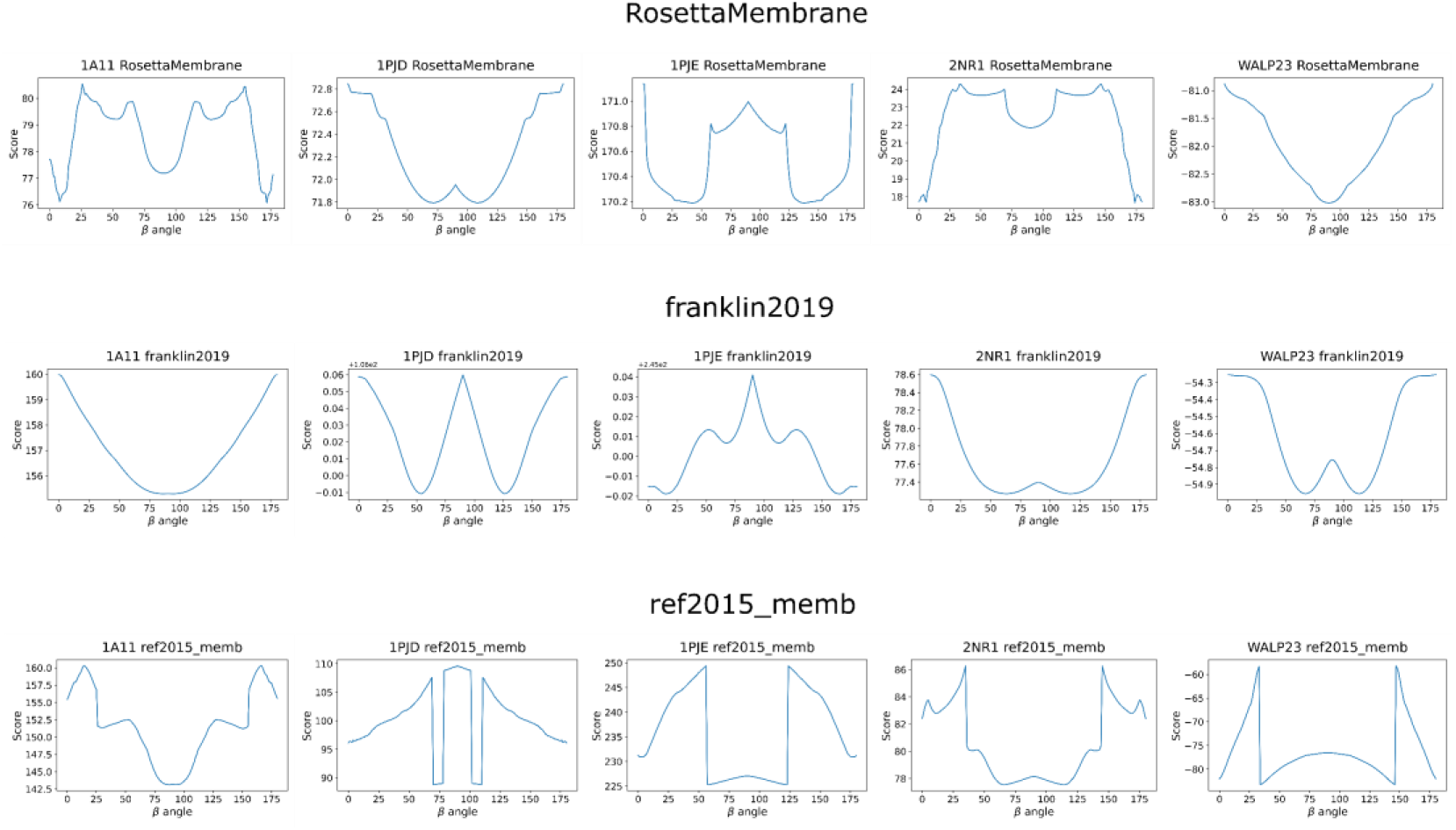
The score versus β-angle (tilt angle) plots calculated for the five hydrophobic peptides by the *RosettaMembrane, franklin2019*, and *ref2015_memb* score functions.

**Supporting Figure 3:**
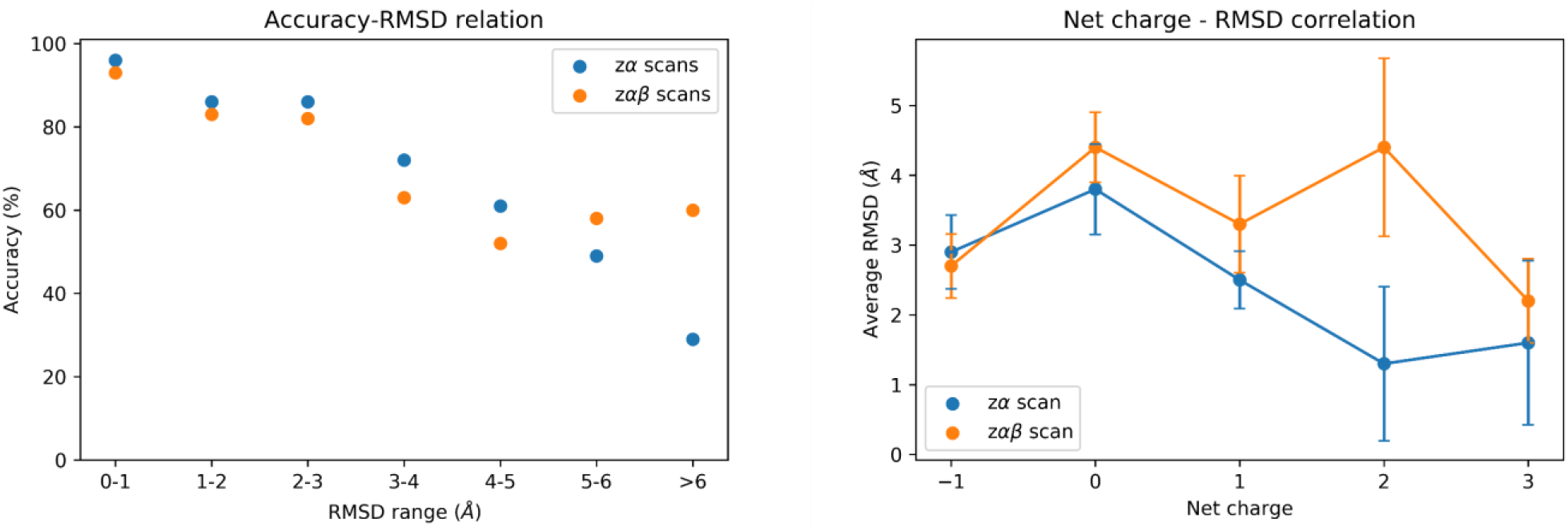
Left panel: The correlation between the calculated RMSD values and the membrane-embedded residue accuracies for the helices in the benchmark set. The accuracy values within 1Å RMSD range were averaged for a clearer representation. Right panel: The correlation between the net charge and the averaged RMSD values bearing the same net charge. Blue represents the result of the zα-scans and orange represents the results of the zαβ-scans.

**Supporting Figure 4:**
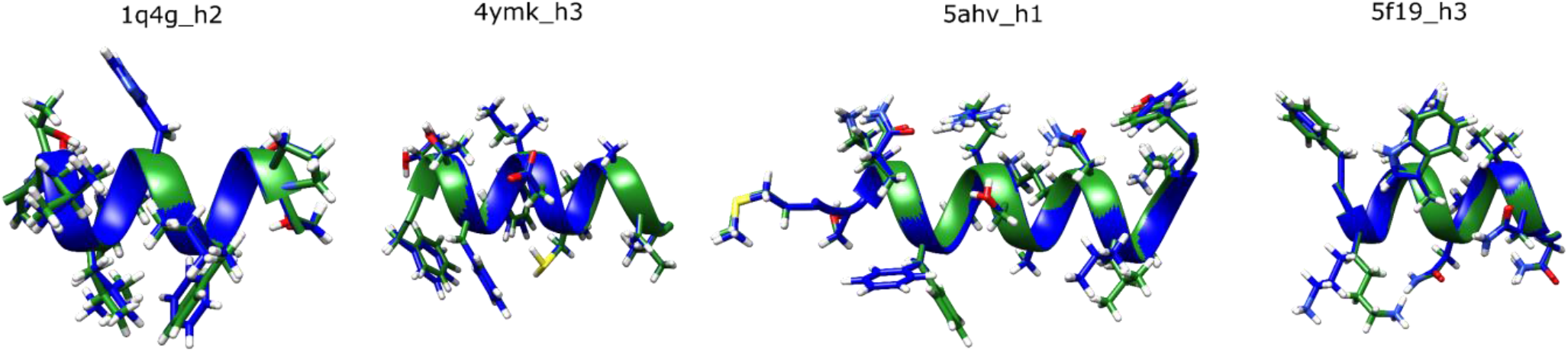
Example structures of the lowest-scoring poses from the calculations with side chain repacking with χ1 recovery rates over 0.85 aligned with the native structures. Green color indicates the native structure and blue color indicates the lowest-scoring Rosetta structure.

